# What can cold-induced transcriptomes of Arctic Brassicaceae tell us about the evolution of cold tolerance?

**DOI:** 10.1101/2021.12.04.471218

**Authors:** Siri Birkeland, Tanja Slotte, Anne K. Brysting, A. Lovisa S. Gustafsson, Torgeir R. Hvidsten, Christian Brochmann, Michael D. Nowak

## Abstract

- Little is known about the evolution of cold tolerance in polar plant species and how they differ from their temperate relatives. To gain insight into their biology and the evolution of cold tolerance, we compared the molecular basis of cold response in three Arctic Brassicaceae species.
- We conducted a comparative time series experiment to examine transcriptional responses to low temperature. RNA was sampled at 22 °C, and after 3h, 6h, and 24h at 2 °C. We then identified sets of genes that were differentially expressed in response to cold and compared them between species, as well as to published data from the temperate *Arabidopsis thaliana*.
- Most differentially expressed genes were species-specific, but a significant portion of the cold response was also shared among species. Among thousands of differentially expressed genes, ∼200 were shared among the three Arctic species and *A. thaliana*, while ∼100 were exclusively shared among the three Arctic species.
- Our results show that cold response differs markedly between Arctic Brassicaceae species, but likely builds on a conserved basis found across the family. They also confirm that highly polygenic traits such as cold tolerance may show little repeatability in their patterns of adaptation.

## Introduction

Only a few plant species have been able to occupy the coldest areas towards the poles (Billings & Mooney, 1968; Wiens & Donoghue, 2004). Transitions from tropical to temperate climates with concurrent development of cold tolerance have been key events in plant evolution (Lancaster & Humphreys, 2020). However, little is known about the evolutionary changes required for a further transition from the temperate to the more extreme polar zones. It is unclear whether polar plant species possess a cold response distinct from that of temperate relatives, and whether it has evolved in a similar, convergent fashion in different species because of similar selection pressures. In general, polar summers are both cooler and shorter than in temperate areas but may resemble certain temperate-alpine summers in terms of climate (Billings, 1974). Polar areas have a unique light environment with up to 24h of daylight in summer, which is important for timing of plant development and circadian rhythm. In the Arctic, the average temperature of the warmest summer month is not more than 10 °C (Elvebakk, 1999), and plants must always be prepared for occasional frost. By studying how Arctic plants cope with low temperatures, we can gain insights into how plants of temperate origins acquire additional cold tolerance, and if there are general trends in plant adaptation to extreme polar environments.

Plants face many challenges upon the transition to colder environments. Low temperatures can affect nearly all aspects of plant cell biochemistry, including protein properties, photosynthesis reactions, cell membrane fluidity, and the creation of gametes (Bomblies *et al*., 2015; Shi *et al*., 2018). Ice formation comes with its own set of challenges and is deadly if ice forms within the cell (Körner, 2003). Some plants can tolerate freezing of the apoplast (the space between the cells), but this may draw water out of the cell and lead to severe dehydration, as well as increase the level of salts and toxic solutes (Steponkus, 1984; Wisniewski & Fuller, 1999; Körner, 2003; Wisniewski *et al*., 2004). In temperate areas, from where most Arctic plants likely are derived (Abbott & Brochmann, 2003), plants prepare for cold periods via cold acclimation, i.e., increasing freezing tolerance in response to low non-freezing temperatures (Thomashow, 2010). Exposing temperate plants to low temperatures typically results in complete reorganization of the transcriptome, ultimately leading to increased freezing tolerance (Kreps *et al*., 2002; Kilian *et al*., 2007; Thomashow, 2010).

The CBF transcription factors (C-repeat-Binding Factors) are among the main “regulatory hubs” of the plant cold response, and have been isolated in many different plant species (Thomashow, 2010; Park *et al*., 2015; Shi *et al*., 2018). The CBFs are induced shortly after exposure to cold stress, and control the expression of >100 cold-regulated (COR) genes downstream (the CBF pathway; Park *et al*., 2015). *Arabidopsis thaliana* exhibits three cold-induced CBFs (*CBF1, CBF2* and *CBF3*, also called *DREB1b, DREB1c* and *DREB1a*; Jia *et al*., 2016), but these transcription factors are not always found in other species (Zhao *et al*., 2012). It is becoming increasingly clear that the CBF regulon involves extensive co-regulation by other, lesser known transcription factors, and that the low temperature regulatory network is complex (Park *et al*., 2015).

One could envision that polar plant species are in less need of a cold acclimation period than temperate species, as temperatures are low year-round and the risk of sudden summer frost is high (e.g., their transcriptomes could be less responsive to a drop in temperature), or that their cold response is somehow more complex (e.g., involving more genes) or more effective (e.g., faster, or involving fewer genes). These hypotheses are relatively unexplored, as there are few in-depth studies of cold-induced transcriptomes in polar plants. Archambault and Strömvik (2011) have studied the Arctic *Oxytropis*, Wang *et al*., (2017) the temperate-subarctic *Eutrema* (*Thellungiella*) *salsugineum*, and Lee *et al*., (2013) the Antarctic *Deschampsia antarctica*. These studies give valuable species-specific information on cold response, but only limited insight into how polar plant species differ from temperate relatives. In this study, we therefore perform a whole-transcriptome investigation of cold response in plant species independently adapted to the high Arctic and compare their response to previously published data from a temperate relative.

The focal species selected for this study, *Cardamine bellidifolia, Cochlearia groenlandica*, and *Draba nivalis*, belong to different temperate lineages that colonized the Arctic independently (Carlsen *et al*., 2009; Jordon-Thaden *et al*., 2010; Koch, 2012) and represent three of the main clades of the Brassicaceae family (clade A, B, and C; divergence time ∼30 Mya; Huang *et al*., 2016). They are ideal model species for studying cold response of Arctic plants for four main reasons: 1) all have their main distribution above the Arctic Circle, 2) they are found in all Arctic bioclimatic subzones (even in polar desert; Elven *et al*., 2011), 3) they belong to the family in which cold response has been most extensively studied, as it includes the model species *A. thaliana* and various economically important crop species (Kilian *et al*., 2007; Park *et al*., 2015; Shi *et al*., 2018), and 4) we recently found evidence of positive selection on genes associated with cold stress in all three species (Birkeland *et al*., 2020). Different genes belonging to similar stress response pathways seem to be under positive selection in different species, suggesting that the three species represent independent adaptation to the Arctic (Birkeland *et al*., 2020). However, our previous study was limited to protein coding regions, and theory predicts that there could be a higher chance of convergence in their expression profiles (Stern, 2013; e.g., Sackton *et al*., 2019). The reason is that mutations in *cis*-regulatory regions should have fewer pleiotropic effects than mutations in coding regions, as protein function may be affected only in a subset of the full expression domain (Gompel & Prud’homme, 2009; Stern, 2013). This is tied to the fact that a gene can have several *cis*-domains and that one *cis*-domain may bind several different transcription factors.

To assess the degree of similarity between Arctic Brassicaceae expression profiles, we subjected the three species to a simulated summer cold shock and identified differentially expressed genes (DEGs) after 3h, 6h and 24h with cold treatment. We aimed to i) characterize the cold-induced transcriptomes of *C. bellidifolia, C. groenlandica*, and *D. nivalis*, ii) describe how their cold response differs from that of their temperate relative *A. thaliana*, and iii) identify potential convergent expression patterns in the three Arctic species.

## Material and methods

### Plant material

For each species, we sowed seeds from a single selfed parent derived from wild Arctic populations from Alaska (Table S1). Several seeds were sown in each of six pots per species, and each pot (with several germinated seeds) was treated as one replicate. Because the plants were siblings derived from selfed parents, and because selfing is assumed to be the predominant mode of reproduction in these species in the wild (Brochmann & Steen, 1999), we expected the six replicates to be close to genetically identical. The plants were grown in a natural daylight room in the phytotron at the University of Oslo from 22^nd^ of March to 18^th^ of May 2018 with day temperature at 22 °C and night temperature at 18 °C. Sunrise was 06:13 March 22nd and 04:33 May 18th, and sunset was 18:36 March 22nd and 21:55 May 18th. Supplementary artificial light was given from 08:00-24:00 to mimic Arctic light conditions (400 W high-intensity discharge lamps), and moisture was at ∼65 % RH.

### Cold shock treatment

Eight weeks after sowing, the plants were given a 24-hour cold shock to simulate a sudden drop in temperature during a typical Arctic summer. At 1:00 p.m. (to minimize correlation with circadian change), we transferred the pots from 22 °C in the daylight room to 2 °C in a cooling room with artificial light from 250 W high-intensity discharge lamps (140-160 µmol m^-2^ s^-1^ measured at plant height) to mimic light conditions in the room where the plants were raised. Leaf tissue was sampled at four time points; just before they were transferred (0h; control), and after 3h, 6h, and 24h. We sampled all six pots for RNA extractions at each time point, but only used the four best RNA extracts per time point for sequencing (i.e., in terms of RNA quality and quantity). These were used as four biological replicates per time point per species. RNA was immediately extracted from fresh, fully expanded leaves as described below.

### RNA extraction and sequencing

For extraction of total RNA, we used the Ambion RNAqueous Kit (Thermo Fisher Scientific, Waltham, USA), following the manufacturer’s protocol for fresh plant tissue: ∼50 mg leaf tissue was immediately ground in Lysis/Binding Solution together with 1 volume of Plant RNA Isolation Aid. The RNA quantity was measured with Broad Range RNA Kit on a Qubit v.2.0 fluorometer (Life Technologies, Carlsbad, USA); RNA quality with an Experion Automated Electrophoresis System Station (Bio-Rad Laboratories, Hercules, USA) and a Nanodrop One spectrophotometer (Thermo Fisher Scientific, Waltham, USA). The Norwegian Sequencing Centre (www.sequencing.uio.no) prepared the libraries using the TruSeq protocol for stranded mRNA (Illumina, San Diego, USA) and performed the sequencing. Samples were indexed, pooled, and run on three lanes (16 samples/lane) on an Illumina HiSeq 3000 (Illumina, San Diego, USA), producing paired end reads with a default insert size of 350 bp and read lengths of 150 bp. The raw reads were quality-checked with FastQC v.0.11.8 (Andrews, 2010), and a single FastQC report per species was generated with MultiQC v.1.7 (Ewels *et al*., 2016).

### Transcriptome assembly and annotation

As there were no available genome assemblies at the start of this study, we assembled a reference transcriptome *de novo* for each species using Trinity v.2.8.5 (Grabherr *et al*., 2011) based on all acquired RNA samples. Trinity was run with the integrated Trimmomatic option (Bolger *et al*., 2014), strand-specificity, and a minimum assembled contig length of 300 bp. The transcriptomes were filtered and annotated with EnTAP (Eukaryotic Non-Model Transcriptome Annotation Pipeline; Hart *et al*., 2020) in two rounds: first to apply the EnTAP filtering option on the raw transcriptome (in order to reduce inflated transcript estimates), and then to annotate the highest expressed isoform and filter out contaminants (used for the annotation of DEGs; see below). For expression filtering, an alignment file was generated with bowtie2 (Langmead & Salzberg, 2012) in combination with RSEM (Li & Dewey, 2011) using default options in the “align_and_estimate_abundance.pl” script provided with the Trinity software suite. Numbers of complete and fragmented BUSCOs (Benchmarking Universal Single-Copy Orthologs) in the filtered transcriptomes were estimated with BUSCO v4.0.6 (Simão *et al*., 2015). The filtered transcriptomes were used as the final reference in the differential expression analyses.

### Differential expression analyses

The Trimmomatic filtered reads were mapped to the reference transcriptomes using the alignment free mapper Salmon with a GC content bias correction (Patro *et al*., 2017). Genes that were differentially expressed after 3h, 6h and 24h of cold treatment were identified with DESeq2 v.1.22.1 (Love *et al*., 2014), using a design formula controlling for the effect of pot number (design = ∼ pot number + time). This means that we tested for the effect of time with 2 °C treatment, while controlling for the individual effects of the sampled pots. A generalized linear model was fitted to each gene and a Wald test (Love *et al*., 2014) applied to test if the 3h, 6h and 24h model coefficients differed significantly from zero when contrasted to the 0h model coefficient. A gene was considered as differentially expressed if the transcript level exhibited ≥ twofold change in response to the cold treatment at the different time points (log2 fold change ≥ 1). P-values were adjusted using the Benjamini-Hochberg procedure (Benjamini & Hochberg, 1995) to control the False Discovery Rate (FDR) and we considered FDR 0.05 as a cutoff for significance. Heatmaps of the top 30 differentially expressed genes with the lowest false discovery rate were generated with the pheatmap package in R (Kolde, 2019) using the regularized log function (rld) on original count data. The mean expression value of a gene was subtracted from each observation prior to heatmap generation.

### Comparison of DEG sets among Arctic species and *A. thaliana*

To enable comparison of DEGs among the Arctic species, and among the Arctic species and *A. thaliana*, we used published data on DEGs in wild type *A. thaliana* in response to 24h chilling treatment (Table S1 in Park *et al*., 2015). In this experiment, *A. thaliana* wild type plants were grown at 22 °C and constant illumination, then exposed to a 4 °C chilling treatment for 24h (Ws-2; see Park *et al*., 2015 for details). We used two different approaches to compare the 24h DEG sets among species. First, we ran OrthoFinder v.2.3.12 (Emms & Kelly, 2019) to identify groups of orthologs among the four species (orthogroups), using the assembled transcriptomes (filtered based on highest expressed isoform) and the Araport11 peptide file of *A. thaliana* downloaded from www.arabidopsis.org. Second, we used the BLASTX (protein-protein) search of BLAST+ v.2.9.0 (Camacho *et al*., 2009) to identify putative *A. thaliana* homologs in the three Arctic species, using the Araport11 peptide file as database and each of the Arctic transcriptome files as query (e-value < 1e-10; choosing the output with the highest bit score). This second approach enabled us to compare gene identity and function more closely among species based on gene pairs. The significance of the overlaps among differentially expressed orthogroups at 24h was evaluated using the supertest function in SuperExactTest v.1.0.7 (Wang *et al*., 2015). We also visualized potential unique overlaps among differentially expressed orthogroups at 24h and among differentially expressed *A. thaliana* homologs at 24h, using UpSetR v.1.4.0 (Conway *et al*., 2017). To compare transcription factor composition, we annotated the *A. thaliana* 24h DEG set with EnTAP (as above).

### Gene function enrichment analyses

To functionally characterize sets of upregulated and downregulated genes, we performed gene ontology (GO) enrichment analyses within the Biological Process (BP), Cellular Component (CC) and Molecular Function (MF) domains for each species and time point. We used Fisher’s exact test in combination with the elim algorithm implemented in topGO 2.34.0 of Bioconductor (Gentleman *et al*., 2004; Alexa *et al*., 2006). The elim algorithm works by traversing the GO-graph bottom-up and discarding genes that already have been annotated to significant child terms, and this is the recommended algorithm by the creators of topGO due to its simplicity (Alexa *et al*., 2006). For the enrichment analyses, we used the gene annotations of the transcriptomes as background gene sets in each test (using the GO-annotations acquired with EnTAP). For *A. thaliana*, we used the org.At.tair.db R package v.3.7.0 to annotate the 24h DEG set of Park *et al*., (2015), and for creating a background gene set used in the enrichment tests (Carlson, 2018). A GO-term was considered significantly enriched if p < 0.05. We did not correct for multiple testing as the enrichment-tests were not independent. Overlaps among enriched GO-terms in similar DEG sets (i.e., upregulated, and downregulated genes at similar time points) were compared among species using UpSetR as above.

### Comparison of DEGs with data set on positively selected genes

In a previous paper, we identified convergent substitutions and tested for positive selection in the same three Arctic Brassicaceae species (see Birkeland *et al*., 2020 for details). We were thus able to check for potential overlaps between positively selected genes/convergent genes and the 24h DEG sets. We blasted the newly assembled transcriptomes against the transcriptomes of our previous study using BLASTX with an e-value cutoff of < 1e-10 and choosing the result with the highest bit score.

### Gene co-expression network analyses

To identify co-expressed gene modules, we performed weighted correlation network analysis for each species using the R package WGCNA (Langfelder and Horvath, 2008). The gene expression matrix was prepared by first filtering out genes with consistent low counts (row sums ≤ 10), and then applying a variance stabilizing transformation within DESeq2. We also filtered out genes with low expression variance by only maintaining genes with variance ranked above the 25 percentile in each data set. A gene adjacency matrix was constructed using default settings and raised to a soft thresholding power of 18 (signed network type). The scale-free topology model fit was only 0.70 for *C. bellidifolia*, 0.79 for *C. groenlandica*, and 0.56 for *D. nivalis* (Fig. S1-S3). However, this is not uncommon when subsets of samples that are globally different from the rest (for instance cold treated versus non-cold treated samples) lead to high correlation among large groups of genes (see Langfelder & Horvath, 2017). The adjacency matrix was translated into a Topological Overlap Matrix (TOM) and co-expressed gene modules were identified by average linkage hierarchical clustering and the Dynamic Tree-Cut algorithm (minimum module size of 30, deepSplit of 2, and pamRespectsDendro as FALSE; Langfelder *et al*., 2008). Closely adjacent modules were merged based on the correlation of their eigengenes (correlation threshold of 0.75). We identified hub genes within each module as the genes with the top 10 % eigengene-based connectivity (module membership). We also calculated the correlation between each module eigengene and a binary cold trait (i.e., cold treated vs. non treated; 1/0) as well as a time-based trait (0h, 3h, 6h, and 24h with cold). Gene function enrichment analyses for each module were performed with topGO as described above. To check for modules overlapping between species, we made UpSet plots based on orthogroup identity. For these plots we included only modules that had a positive correlation ≥0.70 with the binary cold trait or 3h, 6h, or 24h with cold.

## Results

### Transcriptome assemblies and differentially expressed genes (DEGs)

The three *de novo* assemblies contained ∼22,000-24,000 (Trinity) genes and were highly complete in terms of BUSCOs (>90 % complete; Table 1). We identified a gradual increase in the number of DEGs with time at 2 °C from about 700-1000 DEGs after 3h to ∼2500-3000 DEGs after 24h (Table 2; Tables S2-S4). After 3h at 2 °C, most DEGs were upregulated, but after 24h we found similar numbers of downregulated and upregulated DEGs in all species (Table 2). Many 3h and 6h DEGs were also found in the 24h set, but some DEGs were unique for each time point (Tables S5-S7).

**Table 1.** Statistics for de novo transcriptome assemblies.

| Species | Total no. read pairs | No. of “genes” in final assembly<sup>b</sup><br>[raw assembly] | No. of isoforms in final assembly<br>[raw assembly] | % complete BUSCOs in final assembly |
| --- | --- | --- | --- | --- |
| *Cardamine bellidifolia* | 403,256,653<br>(16 samples<sup>a</sup>) | 21,818<br>[42,151] | 42,646<br>[98,419] | 93.8% |
| *Cochlearia groenlandica* | 389,172,001<br>(16 samples<sup>a</sup>) | 22,396<br>[49,768] | 40,639<br>[102,855] | 93.4% |
| *Draba nivalis* | 368,925,923<br>(16 samples<sup>a</sup>) | 23,871<br>[52,096] | 46,282<br>[109,658] | 92.6% |

<sup>a</sup>Four replicates at four time points (0h, 3h, 6h, 24h), <sup>b</sup>Corresponding to Trinity genes (or transcript clusters)

**Table 2.** Number [percentage] of differentially expressed genes (DEGs) after 3h, 6h and 24h with 2 °C. (**bold** = total number of DEGs, ↑/↓ = upregulated/downregulated DEGs)

|  | 3h ✱ | 6h ✱ | 24h ✱ |
| --- | --- | --- | --- |
| *Cardamine bellidifolia* | **1012**<br>857↑, 155↓<br>[85% ↑, 15% ↓] | **1045**<br>877↑, 168↓<br>[84% ↑, 16% ↓] | **2520**<br>(1301↑, 1219↓)<br>[52% ↑, 48% ↓] |
| *Cochlearia groenlandica* | **733**<br>521↑, 212↓<br>[71% ↑, 29% ↓] | **1016**<br>(636↑, 380↓)<br>[63% ↑, 37% ↓] | **3010**<br>(1534↑, 1476↓)<br>[51% ↑, 49% ↓] |
| *Draba nivalis* | **688**<br>(505↑, 183↓)<br>[73% ↑, 27% ↓] | **1583**<br>(998↑, 585↓)<br>[63% ↑, 37% ↓] | **2839**<br>(1484↑, 1355↓)<br>[52% ↑, 48% ↓] |

Most 24h DEGs were species-specific, meaning that they did not belong to shared orthogroups. However, 212 DEGs were shared by the three Arctic species and *A. thaliana*, and 106 were shared by the Arctic species but not by *A. thaliana* (based on orthogroup membership; Fig. 1). When testing for significant overlaps between the four 24h DEG sets, we found that all species combinations had larger overlaps than expected by chance (based on orthogroup identity; all p < 0.01 based on the supertest; Table S8). Thus, our main findings were that 1) the shared portion of the cold response was relatively small, but larger than expected by chance, and 2) more cold responsive genes were shared by the three Arctic species and *A. thaliana* than by the Arctic species alone.

**Fig. 1.**
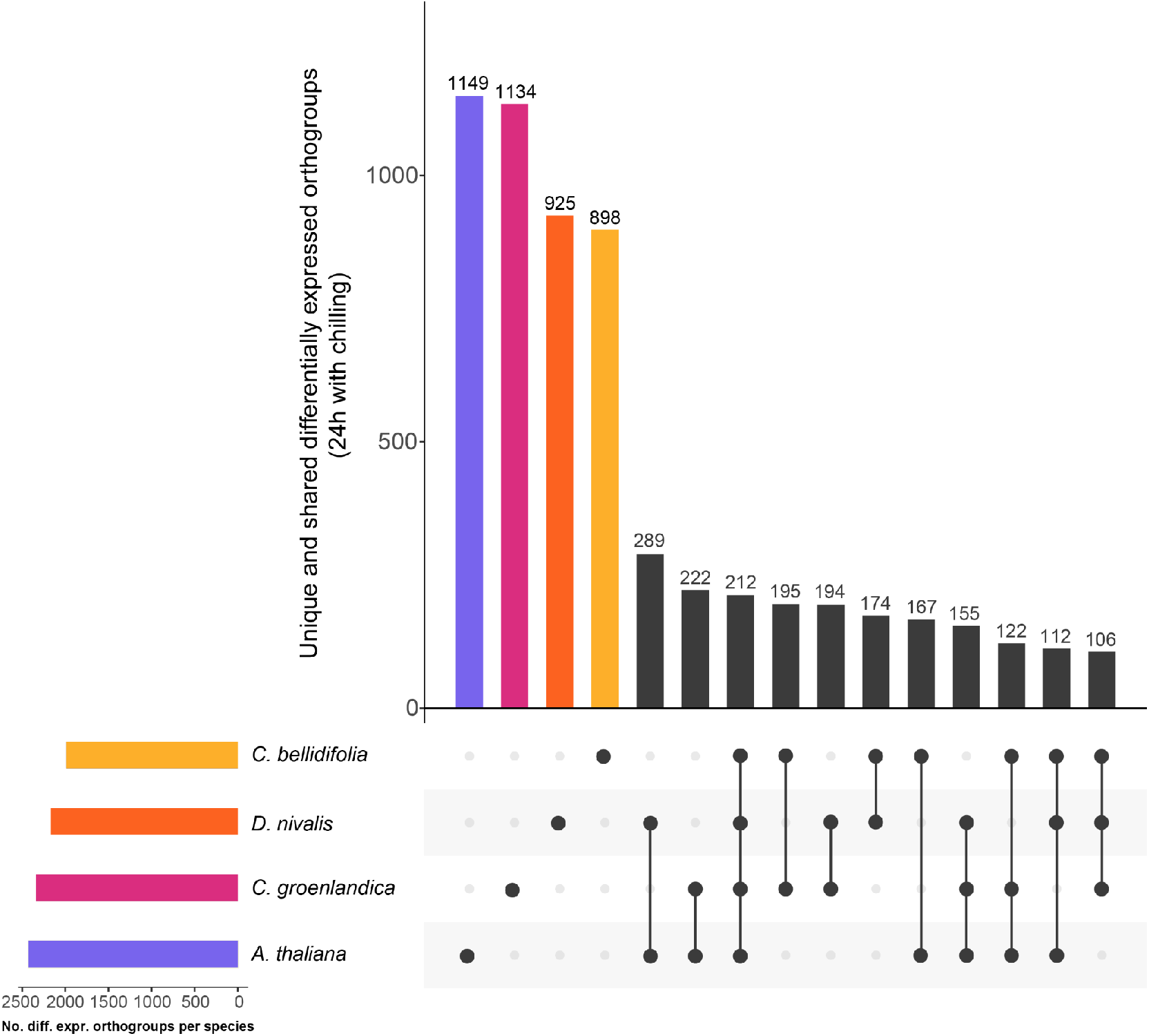
UpSet plot of differentially expressed orthogroups. The plot in the left corner shows total numbers of differentially expressed orthogroups, and the main plot shows the number of unique differentially expressed orthogroups, followed by orthogroup intersections/overlaps between species (connected dots).

### Comparison of DEG annotations among species

Transcription factors made up 9-14% of all Arctic DEGs (Table 3, Tables S2-S4). AP2/ERF was the most common family of transcription factors in the 24h DEG set (Fig. 2); known to include important candidates for cold regulation such as CBFs/DREBs and RAVs. *CBF1* and *CBF4* were upregulated in all Arctic species, and *DREB2A, DREB2C, DREB3*, and *RAV1* were upregulated in one or two Arctic species each (Tables S2-S4). Based on their presence in the DEG sets, other families of transcription factors possibly important for cold response were those containing SANT/Myb domains, MYC-type basic helix-loop-helix (bHLH) domains, basic-leucine zipper domains, and NAC domains (Fig. 2).

**Table 3.** Number of differentially expressed genes (DEGs) annotated with “regulation of transcription” (GO:0006355) after 3h, 6h and 24h with 2 °C. The percentage of the total DEG set is given in parentheses.

|  | 3h ✱ | 6h ✱ | 24h ✱ |
| --- | --- | --- | --- |
| *Cardamine bellidifolia* | **123**<br>(12%) | **139**<br>(13%) | **266**<br>(11%) |
| *Cochlearia groenlandica* | **106**<br>(14%) | **143**<br>(14%) | **344**<br>(11%) |
| *Draba nivalis* | **92**<br>(13%) | **180**<br>(11%) | **260**<br>(9%) |

**Fig. 2.**
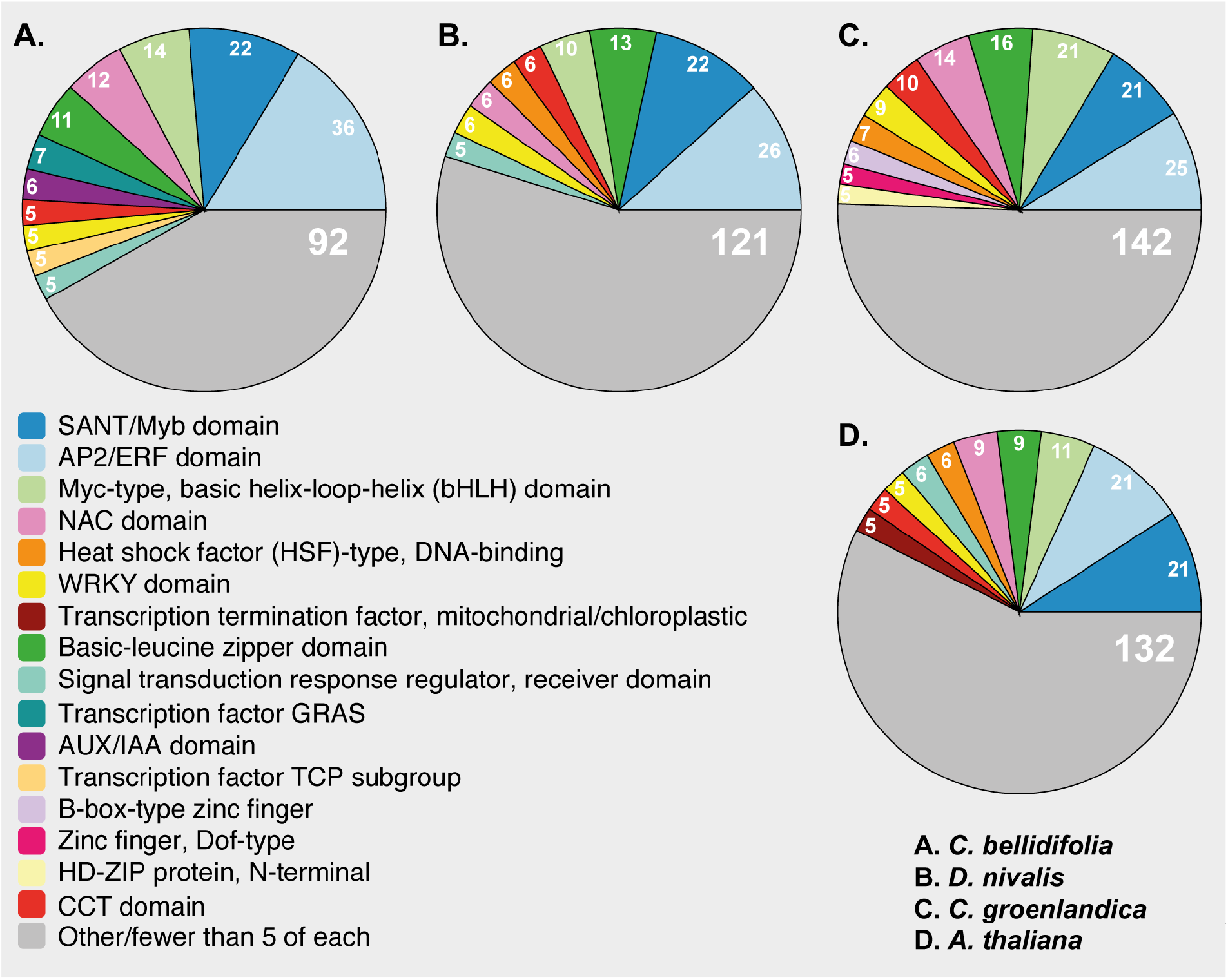
InterPro domains in 24h DEGs annotated with “regulation of transcription” (GO:0006355). Transcription factors that did not have InterPro domain information are not included.

We performed gene function enrichment analyses to functionally characterize the DEG sets and found that most of the significantly enriched GO-terms within the BP, CC, and MF domains were species-specific (Fig. 3, Table 4, Tables S9-S18). However, we found that 20 GO-terms were significantly enriched in all 24h upregulated DEG sets (i.e., in the three Arctic species and *A. thaliana*), while 6 GO-terms were significantly enriched exclusively in the Arctic 24h upregulated DEG sets (BP, CC, and MF; Table S18). Among the GO-terms shared among the Arctic species and *A. thaliana* were many BP terms associated with cold and freezing (e.g., “response to salt stress”, “response to water deprivation”, and “response to oxidative stress”; Figs. 3-4). Shared terms also included those that were associated with the hormones abscisic acid and ethylene, karrikin, and possible cryoprotectants (flavonoids and sucrose; BP, Fig. 4). The GO-terms that were significantly enriched exclusively in the Arctic species included terms such as “spermidine biosynthetic process” (BP) and “arginine decarboxylase activity” (MF). All 24h downregulated DEGs in the Arctic species and *A. thaliana* were enriched for genes associated with growth-related GO-terms such as “phototropism” (BP; all species) and “auxin-activated signaling pathway” (BP; only Arctic species).

**Table 4.**
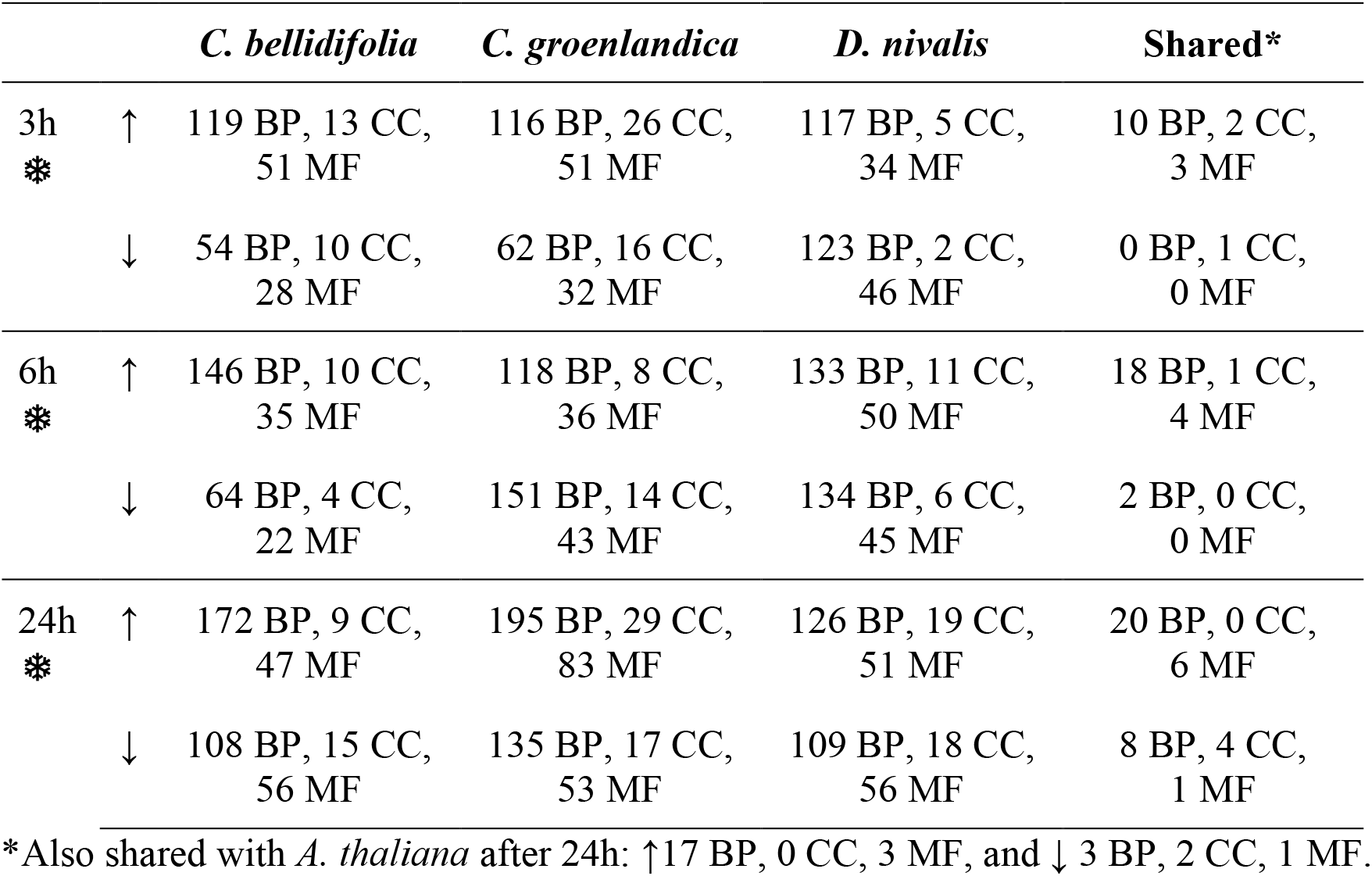
Numbers of significantly enriched GO-terms (p < 0.05) in differentially expressed genes (DEGs) after 3h, 6h and 24h at 2 °C when applying the elim algorithm. Abbreviations: ↑ = Upregulated DEG set, ↓ = Downregulated DEG set, Biological Process = BP, Cellular Component = CC, Molecular Function = MF domains.

**Fig. 3.**
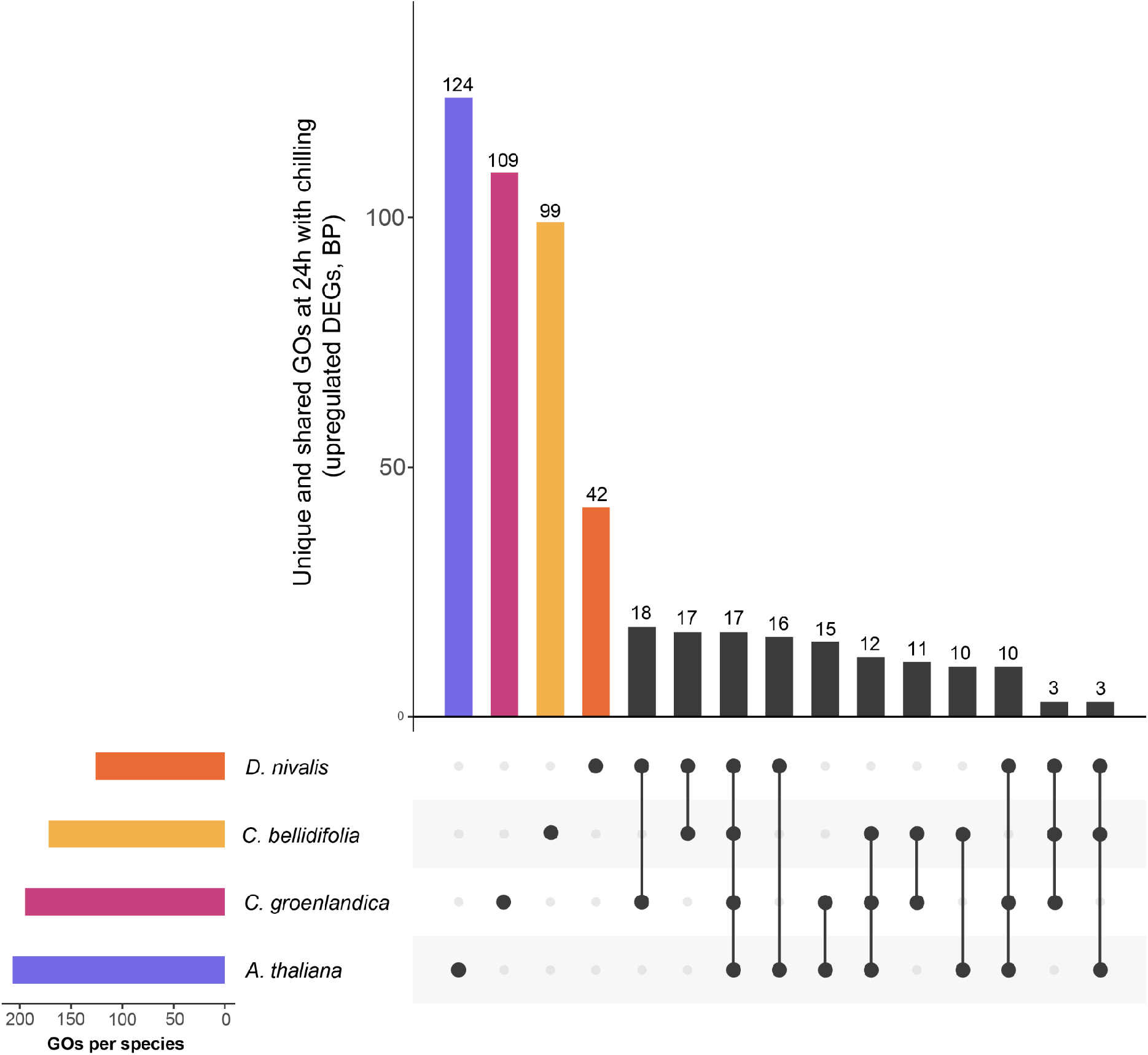
UpSet plot of significantly enriched GO-terms in the 24h upregulated gene sets (Biological Process domain only). The plot in the left corner shows total numbers of significantly enriched GO-terms, and the main plot shows the number of unique significantly enriched GO-terms, followed by the GO-terms that were significantly enriched in more than one species (connected dots).

**Fig. 4.**
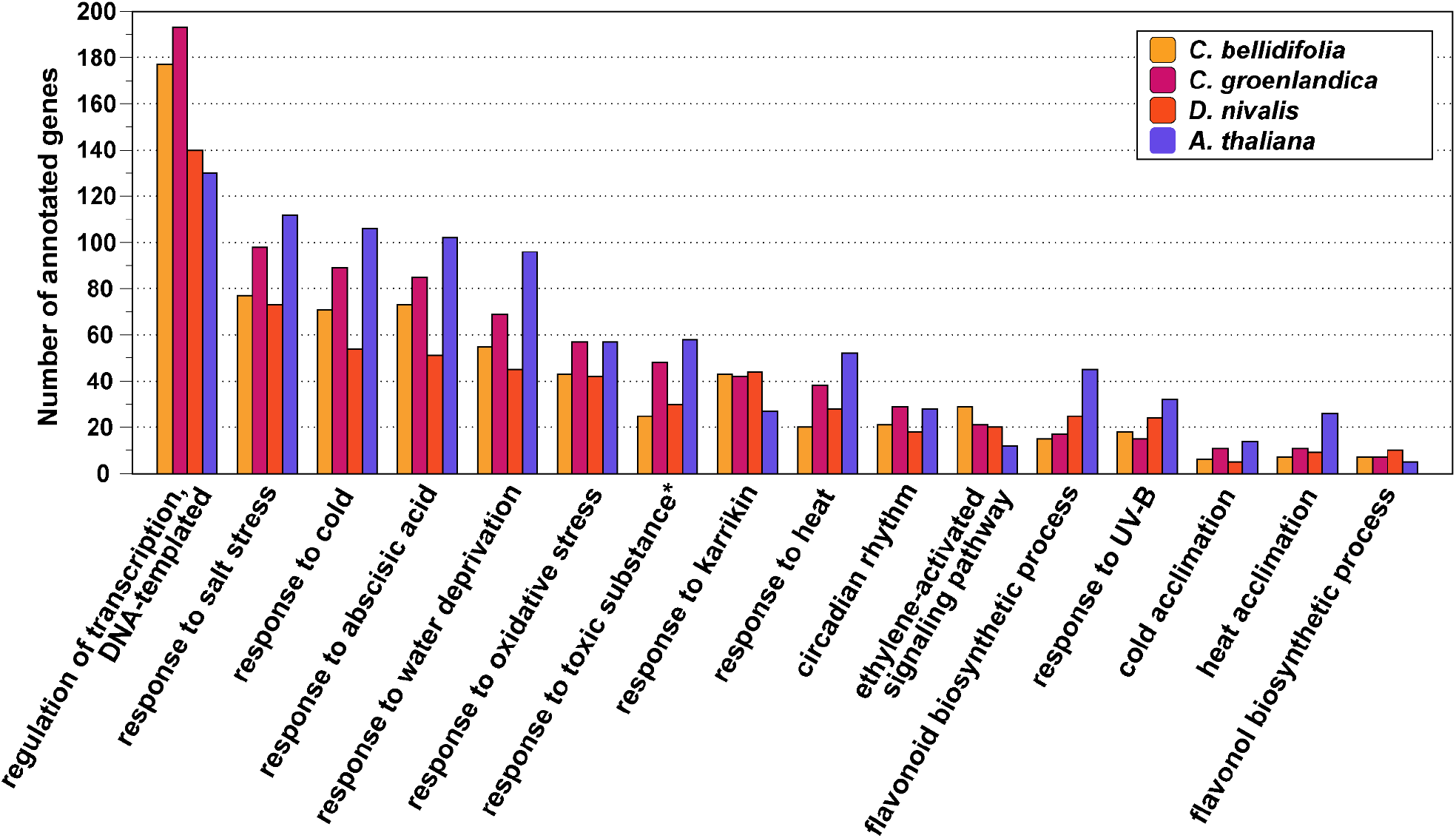
Barchart of genes annotated with Gene Ontology (GO) terms found to be significantly enriched in the 24h upregulated gene set of the three Arctic species (Biological Process domain only). There were 20 overlapping GO-terms among the Arctic 24h upregulated DEG sets, and 17 of these overlapped with *A. thaliana* (BP only). Only terms with at least 10 annotated genes are shown. *Not significantly enriched in *A. thaliana*.

We examined the annotations of the DEGs that were shared exclusively among the three Arctic species, and among the three Arctic species and *A. thaliana* (based on putative *A. thaliana* homologs; Tables S19-S23). The exclusively shared Arctic DEGs included several transcription factors (e.g., *ddf2, ABI5, REVEILLE 2, RAP2*.*2, PCL1*, and *HY5*) and MAPK related genes (*MAPK7, MAPKKK14*, and *MAPKKK18*), while DEGs shared among the Arctic species and *A. thaliana* included known cold induced genes such as *LEA14, COR15B, TCF1, COR27* and *COR28* (upregulated in all species; Tables S19-S20). Many DEGs that were exclusively shared among the Arctic species, as well as among the Arctic species and *A. thaliana*, were found among the topmost differentially expressed genes with the lowest false discovery rate in the Arctic focal species (Fig. S4).

Finally, we also identified 25, 40, and 65 positively selected genes in the DEGs of *C. groenlandica, C. bellidifolia*, and *D. nivalis*, respectively, by blasting our transcriptomes to those of Birkeland *et al*., (2020; Tables S24-S28). Through the same approach we also identified several cold responsive genes previously found to contain convergent substitutions between C. *bellidifolia* and *C. groenlandica, C. bellidifolia* and *D. nivalis*, and/or *C. groenlandica* and *D. nivalis* (Tables S25-S28; supplementary text 1).

### Gene co-expression modules

The gene co-expression network analyses resulted in 13 co-expressed modules in *C. bellidifolia*, 23 in *C. groenlandica* and 14 in *D. nivalis* (after module merging; Fig. 5; Fig. S5-S8). At least one module in each species stood out as highly positively correlated with the binary cold trait (meaning that the eigengene increased continuously with cold; Pearson correlation coefficient > 0.90 and p < 0.001; Fig. 5). These were the light yellow module of *C. bellidifolia*, the light cyan module of *C. groenlandica*, and the dark orange 2 module of *D. nivalis*. They shared 42 co-expressed orthogroups (Fig. S6) and were significantly enriched for 16 of the same GO-terms, including “response to cold”, “response to abscisic acid”, and “regulation of transcription” (Table S29). However, most GO-terms were module or species-specific (Tables S30-S38). The three modules mainly had different hub-genes (i.e., genes with top 10 % highest kME; Tables S39-S41), but *MAPK7* was a hub-gene in the light yellow and dark orange 2 modules (belonging to *C. bellidifolia* and *D. nivalis*, respectively), while *CONSTANS-like 4* was a hub gene in the light cyan and dark orange 2 modules (belonging to *C. groenlandica* and *D. nivalis*, respectively). In addition, several related genes such as *REVEILLE1, 2* and *6*, and *CONSTANS-like 4, 9, 10* and *13* had high kME in all modules, but the exact genes differed among species. The light yellow, light cyan and dark orange 2 modules also included many well-known cold-regulated genes, such as *COR78* and *COR47* (light yellow; *C. bellidifolia*), *CBF1* (light cyan; *C. groenlandica*) and *COR27* (dark orange 2; *D. nivalis*).

**Fig. 5.**
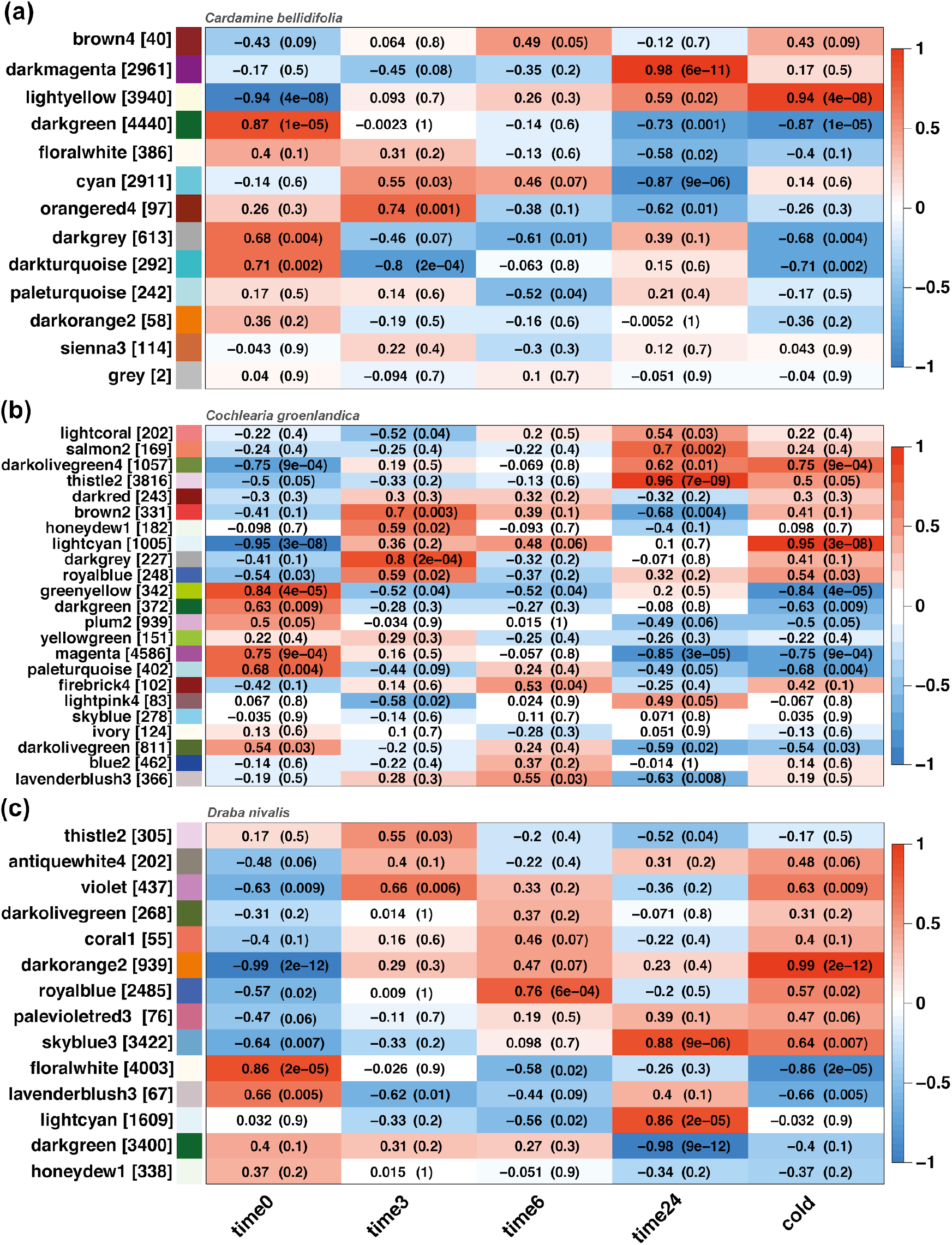
Heatmaps showing Pearson correlation between each co-expression module (module eigengenes) and temperature: 0h, 3h, 6h, and 24h with cold treatment, and a binary measure of cold (cold treatment / no cold treatment). Each row corresponds to a co-expression module and each column to a temperature trait. The number of genes in each module is indicated in brackets after the module name. Cells show the correlation coefficient and corresponding p-value (in parentheses). From the top: a) *Cardamine bellidifolia*, b) *Cochlearia groenlandica* and c) *Draba nivalis*.

Three modules that were positively correlated with 24h of cold stood out as highly overlapping among species (Fig. S7-S8). The dark magenta module in *C. bellidifolia*, the thistle 2 module in *C. groenlandica*, and the sky blue 3 module in *D. nivalis* shared 282 co-expressed orthogroups. In general, the hub genes in these modules varied from species to species, but *RPOT1* was a hub gene in all modules, while *EMB2742, PPC1*, and *GPPS* were hub genes in both dark magenta (*C. bellidifolia*) and skyblue 3 (*D. nivalis*). *MAPK7* (a hub gene in light yellow and dark orange 2 above) was hub-gene in thistle 2 in *C. groenlandica*. The three modules shared 20 significantly enriched GO-terms (domains BP, CC, MF; Table S42), but most GO-terms were module or species-specific (Tables S30-S38). The shared GO-terms were related to metabolism and general cell processes, as well as different cell parts like “nucleus”, “cytosol” and “vacuolar membrane”.

## Discussion

### The cold response in Arctic Brassicaceae is highly species-specific

Our main finding was that the cold response in Arctic Brassicaceae was highly species-specific, but also that the shared portion of the cold response was higher than expected by chance. This suggests that the evolution of cold response in Arctic lineages of this family mainly followed independent genetic trajectories, but with some conserved components. The results broadly agree with our previous study of protein sequence evolution in the same three species, where we found very little overlap in positively selected genes and only a few genes with convergent substitutions (Birkeland *et al*., 2020). We did expect a higher degree of convergence in cold induced expression profiles, but our results agree well with those of a similar study of cold acclimation in the temperate grass subfamily Pooideae where phylogenetically diverse species of grasses showed widespread species-specific transcriptomic responses to low temperatures, but with some conserved aspects (Schubert *et al*., 2019). The proportion of analyzed genes with conserved responses to cold was larger in our study than in that of Schubert *et al*. (16 shared genes with conserved expression profiles), consistent with the shorter divergence time among our species (∼30 Myr in Arctic Brassicaceae vs. ∼65 Myr in Pooideae).

Independent evolutionary trajectories of cold response might be tied to the polygenic nature of this trait, which involves thousands of genes. Highly polygenic traits have high genetic redundancy, and can therefore be expected to show less repeatable patterns of adaptation than traits based on few genes (Yeaman, 2015; Yeaman *et al*., 2016; Barghi *et al*., 2019). Within Brassicaceae, low levels of repeatability have been found in drought tolerance (Marín-de la Rosa *et al*., 2019), which precisely is a trait controlled by many genes. However, there are also studies demonstrating strong convergence in polygenic traits, for example in local adaptation to climate in two distantly related conifers (Yeaman *et al*., 2016), and in flowering time in *Capsella rubella* and *Capsella bursa-pastoris* based on repeated deletions and mutation at the same gene (flowering time is controlled by >60 genes in *A. thaliana*; Slotte *et al*., 2009; Anderson *et al*., 2011; Yang *et al*., 2018). Although it is not known what caused evolutionary repeatability in these two cases, adaptation may end up taking the same routes in the presence of pleiotropic constraints or limited standing genetic variation (as discussed in Gould & Stinchcombe, 2017). Our finding of mainly independent evolution of cold tolerance suggests that such constraints have been of little importance in these Arctic lineages.

### Conserved aspects of the cold response in Brassicaceae

Another major finding in our study was that the cold response in the three Arctic species seems to have more in common with their temperate relative *A. thaliana* than with each other. This shared aspect of the cold response may represent conserved parts of the CBF pathway, which is present also in other plant families (Jaglo *et al*., 2001; Shi *et al*., 2018; Vergara *et al*., 2022). In our study, genes shared between *A. thaliana* and the Arctic species included those tied to circadian regulation and regulation of freezing tolerance (e.g., *COR27* and *COR28*; Li *et al*., 2016). This fits well with the fact that the CBF pathway is gated by the circadian clock (Fowler *et al*., 2005; Dong *et al*., 2011). The continuous increase in DEGs with time at 2 °C also indicates that Arctic plants respond to low temperatures in a way similar to temperate plants (see e.g., Kilian *et al*., 2007; Calixto *et al*., 2018). Thus, this finding suggests Arctic plants are not completely hard-wired to tolerate low temperatures, but that they need a cold acclimation period to develop cold tolerance.

The cold response of our Arctic species included biological processes that have been demonstrated in the cold response of *A. thaliana*, other Brassicaceae species, and species from other plant families (e.g., Zhao *et al*., 2012; Lee *et al*., 2013; Buti *et al*., 2018), including stress responses associated with low temperatures, salt stress, and water deprivation. Stress response pathways associated with cold have been shown to be partially homologous with those of dehydration and salt tolerance (Ingram & Bartels, 1996; Bartels & Souer, 2003; Shamustakimova *et al*., 2017). In addition, ice formation in the apoplast draws water out of the cells and increases the concentration of salts and toxic solutes (Wisniewski & Fuller, 1999; Körner, 2003), leading to cell dehydration (Shi *et al*., 2018). We also found that low temperatures triggered responses to oxidative stress, which accompanies other abiotic stresses such as cold stress, and especially high light intensity in combination with low temperatures (Heino & Palva, 2004; Kilian *et al*., 2007; Lütz, 2010).

We found that genes associated with the hormones abscisic acid (ABA) and ethylene were upregulated in response to cold in all species. ABA is important in plant stress signaling and has been shown to increase in abundance during cold acclimation (Heino & Palva, 2004; Tuteja, 2007). Exogenous application of ABA even increases freezing tolerance in *A. thaliana* and other plants (Thomashow, 1999). Ethylene is also reported to influence freezing tolerance and regulates the CBF pathway itself (Kazan, 2015). In most plant species, this entails a positive regulation of freezing tolerance (e.g., in tomato and tobacco; Zhang & Huang, 2010), although contradicting results have been found in *A. thaliana* (Kazan, 2015). Thus, ABA and ethylene may have important roles in a shared cold response machinery in the Brassicaceae and possibly in other plant families.

Plants may increase their freezing tolerance by accumulating compounds that hinder ice nucleation or by alleviating the effects of ice formation by protecting plant tissues against freezing damage (i.e. cryoprotectants; Ruelland *et al*., 2009). We observed that genes associated with sucrose and flavonoids (especially flavonols) were upregulated in response to cold in all species. Sucrose is a powerful cryoprotectant that lowers the freezing point and thus leads to supercooling and avoidance of freezing (Reyes-Díaz *et al*., 2006), diminishes the water potential between the apoplastic space and the cell in the face of ice formation (osmotic adjustment; Ruelland *et al*., 2009), and regulates cold acclimation itself (Rekarte-Cowie *et al*., 2008). Accumulation of flavonoids is associated with cold acclimation and higher freezing tolerance (Schulz *et al*., 2016). In particular, flavonols might have a role in protecting cell membranes during freezing stress (Korn *et al*., 2008). In addition, karrikin seems to be important in acquiring cold/freezing tolerance in Brassicaceae. This compound has received little attention in relation to cold stress, but it inhibits germination under unfavorable conditions and increases plant vigor in the face of abiotic stress through regulation of redox homeostasis (Shah *et al*., 2020). Thus flavonoids, sucrose, and karrikin seem to have important roles in the cold response of four distantly related Brassicaceae species.

The shared trends outlined by this study indicate that cold response is built upon a similar scaffold in Brassicaceae and support the claim that cold tolerance is a complex trait that is difficult to evolve (Donoghue, 2008). Our results suggest that the last common ancestor of the core Brassicaceae possessed some kind of cold tolerance machinery. The major clades of the Brassicaceae are thought to have radiated in response to the change to a colder and drier climate at the Eocene-Oligocene transition (∼33.9 Ma; Huang *et al*., 2016), suggesting that evolution of a basic cold tolerance machinery contributed to its successful colonization of cold and dry regions. Nevertheless, the highly species-specific cold response we found in our Arctic species also suggests that there is great evolutionary flexibility in cold coping strategies once there is a basis to build upon.

### The Arctic cold response

The ∼100 genes we identified as exclusively shared among the three Arctic species included many transcription factors that potentially may have large effects. Among these, *ddf2* may be an important candidate for adaptation to cold conditions as it is closely related to *CBF1-3* (genes with key regulatory roles in the CBF pathway). Overexpression of *ddf2* leads to dwarfism and late flowering in *A. thaliana* (the full gene name is ‘dwarf and delayed flowering 2’; Magome *et al*., 2004), which coincides well with the prostate growth that often characterizes Arctic plants. So far, it seems like *ddf2* is not cold regulated in *A. thaliana*, but the sister gene *ddf1* is upregulated in response to salinity stress (Medina *et al*., 2011). Another interesting transcription factor in relation to Arctic cold regulation is *RAP2*.*2*, which is related to *DEAR1* - a transcription factor known to mediate freezing stress responses in plants (Tsutsui *et al*., 2009). We also found several shared transcription factors related to circadian rhythm, such as *REVEILLE 2, PCL1* and *HY5*. We cannot rule out that our experimental light regime may have provoked the expression of these genes, but the interplay between the circadian clock and the CBF pathway is known to be important for balancing freezing tolerance and plant growth (e.g. Dong *et al*., 2011; Shi *et al*., 2018). In addition, *REVEILLE 1, 2* and *6* had high eigengene-based connectivity in cold correlated co-expression modules in all species. One could therefore speculate that the Arctic light regime has triggered the evolution of new links between the CBF pathway and the circadian clock. Our finding of exclusively shared transcription factors among the Arctic species fits well with theory predicting that hub genes may be important hot spots for convergent evolution (Martin & Orgogozo, 2013; Yeaman *et al*., 2016).

The exclusively shared Arctic DEGs further contained traces of a Mitogen Activated Protein Kinase cascade (*MAPK7, MAPKKK14, MAPKKK18*). Such cascades are known to be important in regulating the CBF pathway (Teige *et al*., 2004; Shi *et al*., 2018). A potential role in cold regulation is supported by our finding of *MAPK7* as a hub gene in several co-expression modules highly correlated with cold. In the temperate *A. thaliana*, it has been shown that the *MKK2* pathway regulates cold stress signaling (Teige *et al*., 2004), and that the sister gene *MAPK6* is involved in releasing inhibitory effects on CBF gene expression (Kim *et al*., 2017). The shared MAPK and MAPKKKs may therefore have important roles in Arctic cold responses.

Although we found that the Arctic DEG sets were functionally similar to that of the temperate *A. thaliana*, spermidine related genes were only overrepresented in the Arctic upregulated DEG sets. Spermidine is a polyamine that may be important for maintaining photosynthesis rates during low temperatures as cucumber plants pretreated with spermidine show less decline in photosynthesis rates during chilling than non-treated plants (He *et al*., 2002). Sustaining photosynthesis at low temperatures is especially important for Arctic plants as temperatures are low in summer, and they have optimum photosynthesis rates at lower temperatures than other plants (Chapin, 1983). The significant enrichment we found was caused by only a few cold induced genes related to spermidine. One of these, *ADC1*, which has been found to be involved in acquiring freezing tolerance in wild potato (Kou *et al*., 2018), was also responsible for the significant enrichment of arginine decarboxylase activity found in the Arctic species. This makes it an interesting candidate for possible adaptation to cold conditions in Arctic species.

We also observed that several of the genes we previously identified as being under positive selection or containing convergent substitutions in the Arctic species also are cold responsive (Birkeland *et al*., 2020). These included genes such as *CSDP1* and *COR15B* in *D. nivalis*, two transmembrane proteins in *C. bellidifolia*, and *LEA4-5* in *C. groenlandica*. The convergent *EMB2742* and *MAPKKK14* were upregulated in response to cold in all three species, which indicates that they might have an important function in cold tolerance. Since our previous analyses of positive selection and convergent molecular evolution were not based on cold induced transcriptomes, it is possible that additional positively selected genes may exist among our cold induced DEGs.

### Conclusions, limitations, and future perspectives

We have presented the first comparative study of Arctic cold-induced transcriptomes, providing new insight into the evolution of cold response in the Brassicaceae. We found that the cold response in different species of Arctic Brassicaceae has more in common with their temperate relative *A. thaliana* than they have with each other. Certain transcription factors, a potential *MAPK* cascade, and the ability to perform photosynthesis under low temperatures, may nevertheless represent important shared adaptations to the Arctic biome. Here we only included published data from one temperate species, and it is therefore possible that some of the ∼100 genes we found to be exclusively shared among the Arctic species can be found in other temperate Brassicaceae as well. In any case, our results clearly show that there is not a single, but many ways Arctic plants may respond to low temperatures. Considering the polygenic nature of cold response, and that the three species descend from different temperate relatives, it would perhaps have been more surprising if their cold response had converged into something more uniquely Arctic.

Our comparative approach gave valuable insights into the basis of a conserved cold response possibly found throughout the Brassicaceae family. This shared cold response includes well known genes in the CBF pathway and may be co-regulated by abscisic acid and ethylene. It also included upregulation of genes associated with sucrose, flavonoids, and karrikin. Our results are based on comparisons between our new data for the Arctic species and previously published data for *A. thaliana*, and we note that there might be methodological differences that may affect the number of DEGs considered as significant and thus the number of shared genes. Our *de novo* assembled transcriptomes may contain some inaccuracies in the delimitation of genes (e.g., isoforms of the same gene mistakenly being delineated as different genes). However, we used a stringed filtering scheme to reduce inflated transcript numbers, and such differences should not affect the overall results.

Further studies are needed to verify the expression of cold regulated genes shared among the Arctic species, as well as among *A. thaliana* and the Arctic species, and to obtain a better understanding of species-specific gene function. Such investigations should address the effects of shared transcription factors and MAPK genes on cold and freezing tolerance. This introductory study has provided novel insights into how Arctic plants may respond to a summer cold shock, but more studies are needed to learn how they cope with long-term cold stress, as well as cold stress in combination with the light regime typical of Arctic environments.

## Supporting information

Table S2. DEGs with annotations: A) 3h, B) 6h, C) 24h Cardamine bellidifolia

Table S3. DEGs with annotations: A) 3h, B) 6h, C) 24h Cochlearia groenlandica

Table S4. DEGs with annotations: A) 3h, B) 6h, C) 24h Draba nivalis

Table S9. topGO tables for Cardamine bellidifolia DEGs, Biological Process

Table S10. topGO tables for Cardamine bellidifolia DEGs, Cellular Component

Table S11. topGO tables for Cardamine bellidifolia DEGs, Molecular Function

Table S12. topGO tables for Cochlearia groenlandica DEGs, Biological Process

Table S13. topGO tables for Cochlearia groenlandica DEGs, Cellular Component

Table S14. topGO tables for Cochlearia groenlandica DEGs, Molecular Function

Table S15. topGO tables for Draba nivalis DEGs, Biological Process

Table S16. topGO tables for Draba nivalis DEGs, Cellular Component

Table S17. topGO tables for Draba nivalis DEGs, Molecular Function

Table S19. Genes found only in Arctic 24h DEG sets (based on putative A. thaliana homologs)

Table S20. Genes shared by A. thaliana and Arctic species in 24h DEG sets (based on putative A. thaliana homologs)

Table S21. Cardamine bellidifolia DEGs and putative Arabidopsis thaliana homologs: BLAST results

Table S22. Cochlearia groenlandica DEGs and putative Arabidopsis thaliana homologs: BLAST results

Table S23. Draba nivalis DEGs and putative Arabidopsis thaliana homologs: BLAST results

Table S26. Cardamine bellidifolia DEGs blasted against the Alaskan C. bellidifolia transcriptome of Birkeland et al. 2020

Table S27. Cochlearia groenlandica DEGs blasted against the Alaskan C. groenlandica transcriptome of Birkeland et al. 2020

Table S28. Draba nivalis DEGs blasted against the Alaskan D. nivalis transcriptome of Birkeland et al. 2020

Table S30. topGO tables for Cardamine bellidifolia Co-expression modules, Biological Process

Table S31. topGO tables for Cardamine bellidifolia Co-expression modules, Cellular Component

Table S32. topGO tables for Cardamine bellidifolia Co-expression modules, Molecular Function

Table S33. topGO tables for Cochlearia groenlandica Co-expression modules, Biological Process

Table S34. topGO tables for Cochlearia groenlandica Co-expression modules, Cellular Component

Table S35. topGO tables for Cochlearia groenlandica Co-expression modules, Molecular Function

Table S36. topGO tables for Draba nivalis Co-expression modules, Biological Process

Table S37. topGO tables for Draba nivalis Co-expression modules, Cellular Component

Table S38. topGO tables for Draba nivalis Co-expression modules, Molecular Function

Table S39. Annotated hub genes, Cardamine bellidifolia

Table S40. Annotated hub genes, Cochlearia groenlandica

Table S41. Annotated hub genes, Draba nivalis

## Acknowledgements

We thank The Phytotron at the University of Oslo, and engineers Bjørn Langrekken, Marit Langrekken, and Ingrid Johansen for help with the technical setup for this experiment. We also thank the SpeciationClock Team, Xuyue Yang, Marian Schubert, Aelys Humphreys, Lars Grønvold, William Hughes, and Marco Fracassetti for valuable input and comments. Computational analyses were performed on the Saga Cluster owned by Uninett/Sigma2, as well as the Abel Cluster owned by Uninett/Sigma2 and the University of Oslo (operated by the Department for Research Computing at USIT, the University of Oslo IT-department; http://www.hpc.uio.no/). The study was funded by the Research Council of Norway through the SpArc project (RCN 240223) awarded to C.B., with additions from EVOTREE project (RCN 287465) awarded to T.R.H.

## Author Contributions

MDN conceived the idea for the study. SB, MDN, and TS designed and built the study. ALSG collected the Arctic plant material. SB and MDN carried out the cold shock treatment and experimental plant work for RNA sequencing, with input from ALSG and AKB. SB carried out all downstream analyses with input from MDN, TS, TRH, CB, AKB, and ALSG. SB wrote the manuscript with comments from CB, AKB, ALSG, TS, MDN, and TRH. CB secured the funding for the study with contributions from MDN and TRH.

## Data availability

Full lists of differentially expressed genes, shared differentially expressed genes, gene annotations, and gene function enrichment results are available as Supplementary Material. RNA sequences for the three Arctic species are available at the NCBI Sequence Read Archive (BioSample ID: SAMN27155772-SAMN27155819, BioProject ID: PRJNA821902), while their transcriptome assemblies have been deposited at datadryad.org (https://doi.org/10.5061/dryad.hhmgqnkjh).

