## Supplementary material for "What can cold-induced transcriptomes of Arctic Brassicaceae tell us about the evolution of cold tolerance?"

### Supplementary figures

**Figure S1.** Background information for *Cardamine bellidifolia* WGCNA
**A)** Sample clustering
**B)** Soft Thresholding Powers plotted against Scale Free Topology Fit and Mean Connectivity

**Figure S2.** Background information for *Cochlearia groenlandica* WGCNA
**A)** Sample clustering
**B)** Soft Thresholding Powers plotted against Scale Free Topology Fit and Mean Connectivity

**Figure S3.** Background information for *Draba nivalis* WGCNA
**A)** Sample clustering
**B)** Soft Thresholding Powers plotted against Scale Free Topology Fit and Mean Connectivity

**Figure S4.** Heatmaps of the most significantly differentially expressed genes (DEGs)

**Figure S5.** Dendrograms with module colors (WGCNA)

**Figure S6.** UpSet plot of modules most positively correlated with the binary cold trait

**Figure S7.** UpSet plot of modules most positively correlated with 24h of cold

**Figure S8.** UpSet plots of other modules positively correlated (≥ 0.70) with cold traits
**A)** All modules
**B)** All 24h modules
**C)** All 3h and 6h modules

### Supplementary tables

**Table S1.** Sampling information for Arctic focal species

**Table S2.** DEGs with annotations: **A)** 3h, **B)** 6h, **C)** 24h
*Cardamine bellidifolia*. *Separate excel file*.

**Table S3.** DEGs with annotations: **A)** 3h, **B)** 6h, **C)** 24h
*Cochlearia groenlandica. Separate excel file*.

**Table S4.** DEGs with annotations: **A)** 3h, **B)** 6h, **C)** 24h
*Draba nivalis*. *Separate excel file*.

**Table S5.** Overlapping DEGs among time points in *Cardamine bellidifolia*

**Table S6.** Overlapping DEGs among time points in *Cochlearia groenlandica*

**Table S7.** Overlapping DEGs among time points in *Draba nivalis*

**Table S8.** Results from the supertest.
Intersections of differentially expressed orthogroups.

**Table S9.** topGO tables for *Cardamine bellidifolia* DEGs, Biological Process.
*Separate excel file*.

**A)** upregulated DEGs 3h
**B)** upregulated DEGs 6h
**C)** upregulated DEGs 24h
**D)** downregulated DEGs 3h
**E)** downregulated DEGs 6h
**F)** downregulated DEGs 24h

**Table S10.** topGO tables for *Cardamine bellidifolia* DEGs, Cellular Component.
*Separate excel file.*
**A)** upregulated DEGs 3h
**B)** upregulated DEGs 6h
**C)** upregulated DEGs 24h
**D)** downregulated DEGs 3h
**E)** downregulated DEGs 6h
**F)** downregulated DEGs 24h

**Table S11.** topGO tables for *Cardamine bellidifolia* DEGs, Molecular Function.
*Separate excel file.*
**A)** upregulated DEGs 3h
**B)** upregulated DEGs 6h
**C)** upregulated DEGs 24h
**D)** downregulated DEGs 3h
**E)** downregulated DEGs 6h
**F)** downregulated DEGs 24h

**Table S12.** topGO tables for *Cochlearia groenlandica* DEGs, Biological Process.
*Separate excel file.*
**A)** upregulated DEGs 3h
**B)** upregulated DEGs 6h
**C)** upregulated DEGs 24h
**D)** downregulated DEGs 3h
**E)** downregulated DEGs 6h
**F)** downregulated DEGs 24h

**Table S13.** topGO tables for *Cochlearia groenlandica* DEGs, Cellular Component.
*Separate excel file.*
**A)** upregulated DEGs 3h
**B)** upregulated DEGs 6h
**C)** upregulated DEGs 24h
**D)** downregulated DEGs 3h
**E)** downregulated DEGs 6h
**F)** downregulated DEGs 24h

**Table S14.** topGO tables for *Cochlearia groenlandica* DEGs, Molecular Function.
*Separate excel file.*
**A)** upregulated DEGs 3h
**B)** upregulated DEGs 6h
**C)** upregulated DEGs 24h
**D)** downregulated DEGs 3h
**E)** downregulated DEGs 6h
**F)** downregulated DEGs 24h

**Table S15.** topGO tables for *Draba nivalis* DEGs, Biological Process.
*Separate excel file.*

**A)** upregulated DEGs 3h
**B)** upregulated DEGs 6h
**C)** upregulated DEGs 24h
**D)** downregulated DEGs 3h
**E)** downregulated DEGs 6h
**F)** downregulated DEGs 24h

**Table S16.** topGO tables for *Draba nivalis* DEGs, Cellular Component.
*Separate excel file*.
**A)** upregulated DEGs 3h
**B)** upregulated DEGs 6h
**C)** upregulated DEGs 24h
**D)** downregulated DEGs 3h
**E)** downregulated DEGs 6h
**F)** downregulated DEGs 24h

**Table S17.** topGO tables for *Draba nivalis* DEGs, Molecular Function.
*Separate excel file*.
**A)** upregulated DEGs 3h
**B)** upregulated DEGs 6h
**C)** upregulated DEGs 24h
**D)** downregulated DEGs 3h
**E)** downregulated DEGs 6h
**F)** downregulated DEGs 24h

**Table S18.** Significantly enriched GO-terms shared by all Arctic species (all DEG sets)

**Table S19.** Genes found only in Arctic 24h DEG sets (based on putative *A. thaliana homologs*). *Separate excel file*.

**Table S20.** Genes shared by *A. thaliana* and Arctic species in 24h DEG sets (based on putative *A. thaliana homologs*). *Separate excel file*.

**Table S21.** *Cardamine bellidifolia* DEGs and putative *Arabidopsis thaliana* homologs: BLAST results. *Separate excel file*.
**A)** upregulated DEGs 3h
**B)** upregulated DEGs 6h
**C)** upregulated DEGs 24h
**D)** downregulated DEGs 3h
**E)** downregulated DEGs 6h
**F)** downregulated DEGs 24h

**Table S22.** *Cochlearia groenlandica* DEGs and putative *Arabidopsis thaliana* homologs: BLAST results. *Separate excel file*.
**A)** upregulated DEGs 3h
**B)** upregulated DEGs 6h
**C)** upregulated DEGs 24h
**D)** downregulated DEGs 3h
**E)** downregulated DEGs 6h
**F)** downregulated DEGs 24h

**Table S23.** *Draba nivalis* DEGs and putative *Arabidopsis thaliana* homologs: BLAST results. *Separate excel file*.
**A)** upregulated DEGs 3h

**B)** upregulated DEGs 6h
**C)** upregulated DEGs 24h
**D)** downregulated DEGs 3h
**E)** downregulated DEGs 6h
**F)** downregulated DEGs 24h

**Table S24.** Positively selected genes from Birkeland et al. 2020 found in Arctic DEG sets.

**Table S25.** Convergent genes from Birkeland et al. 2020 found in Arctic DEG sets.

**Table S26.** *Cardamine bellidifolia* DEGs blasted against the Alaskan *C. bellidifolia* transcriptome of Birkeland et al. 2020. *Separate excel file*.
A) Blast results for all up-regulated genes
B) Blast results for all down-regulated genes
C) DEGs under positive selection
D) DEGs with convergent substitutions in Arctic species

**Table S27.** *Cochlearia groenlandica* DEGs blasted against the Alaskan *C. groenlandica* transcriptome of Birkeland et al. 2020. *Separate excel file*.
A) Blast results for all up-regulated genes
B) Blast results for all down-regulated genes
C) DEGs under positive selection
D) DEGs with convergent substitutions in Arctic species

**Table S28.** *Draba nivalis* DEGs blasted against the Alaskan *D. nivalis* transcriptome of Birkeland et al. 2020. *Separate excel file*.
A) Blast results for all up-regulated genes
B) Blast results for all down-regulated genes
C) DEGs under positive selection
D) DEGs with convergent substitutions in Arctic species

**Table S29.** Significantly enriched GO-terms shared by the light yellow (*C. bellidifolia*) light cyan (*C. groenlandica*) and dark orange 2 (*D. nivalis*) co-expression modules.

**Table S30.** topGO tables for *Cardamine bellidifolia* Co-expression modules, Biological Process.
*Separate excel file*.

**Table S31.** topGO tables for *Cardamine bellidifolia* Co-expression modules, Cellular Component.
*Separate excel file*.

**Table S32.** topGO tables for *Cardamine bellidifolia* Co-expression modules, Molecular Function.
*Separate excel file*.

**Table S33.** topGO tables for *Cochlearia groenlandica* Co-expression modules, Biological Process.
*Separate excel file*.

**Table S34.** topGO tables for *Cochlearia groenlandica* Co-expression modules, Cellular Component.
*Separate excel file*.

**Table S35.** topGO tables for *Cochlearia groenlandica* Co-expression modules,

Molecular Function.
*Separate excel file*.

**Table S36.** topGO tables for *Draba nivalis* Co-expression modules, Biological Process. *Separate excel file*.

**Table S37.** topGO tables for *Draba nivalis* Co-expression modules, Cellular Component. *Separate excel file*.

**Table S38.** topGO tables for *Draba nivalis* Co-expression modules, Molecular Function. *Separate excel file*.

**Table S39.** Annotated hub genes, *Cardamine bellidifolia. Separate excel file*.

**Table S40.** Annotated hub genes, *Cochlearia groenlandica. Separate excel file*.

**Table S41.** Annotated hub genes, *Draba nivalis. Separate excel file*.

**Table S42.** Significantly enriched GO-terms shared by the dark magenta (*C. bellidifolia*) thistle 2 (*C. groenlandica*) and sky blue 3 (*D. nivalis*) co-expression modules.

### Supplementary text

**Text S1.** Note on positively selected/convergent cold-responsive genes

**A.**

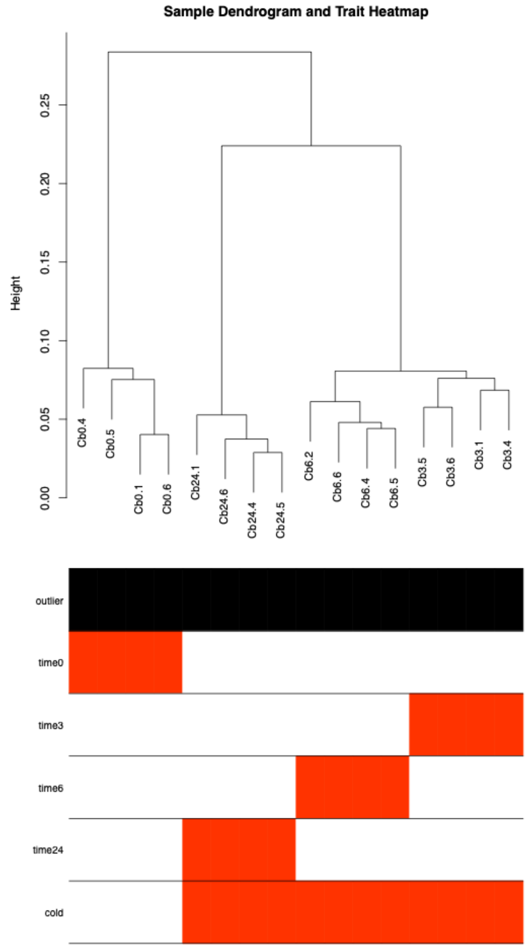


**B.**

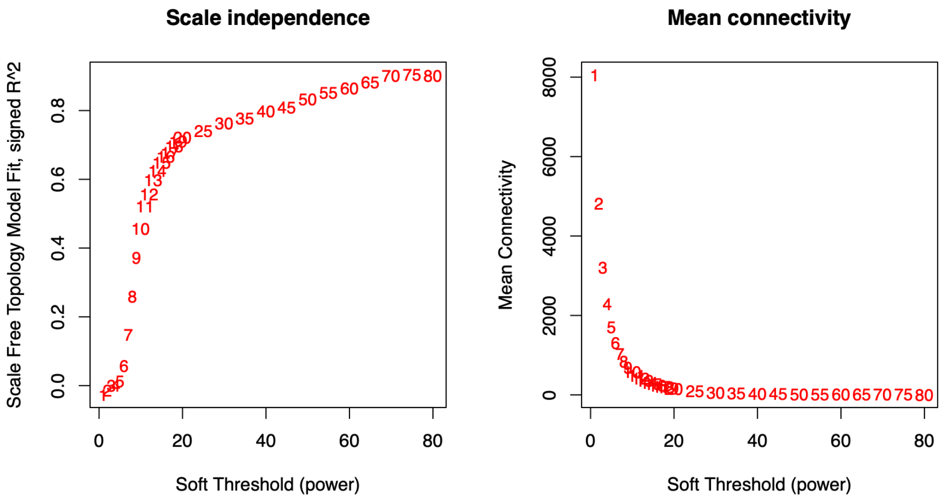


**Supplementary figure 1. Background information for *Cardamine bellidifolia* WGCNA.** A: Plot showing clustering of samples, B: Soft Thresholding Powers plotted against Scale Free Topology Fit (left) and Mean Connectivity (right).

**A.**

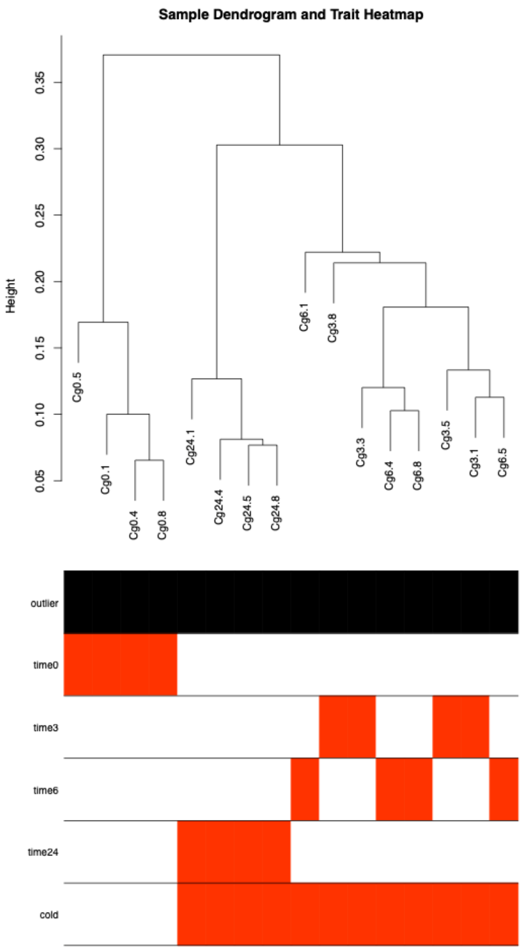


**B.**

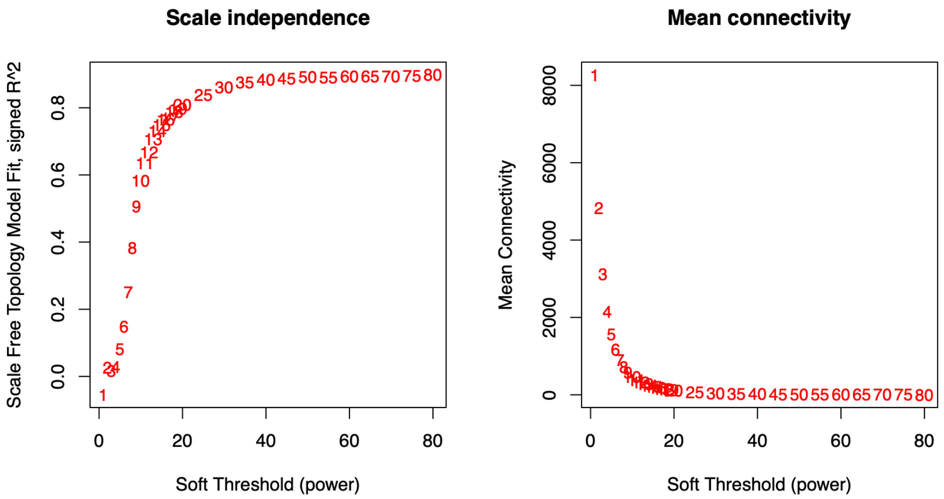


**Supplementary figure 2. Background information for *Cochlearia groenlandica* WGCNA.** A: Plot showing clustering of samples, B: Soft Thresholding Powers plotted against Scale Free Topology Fit (left) and Mean Connectivity (right).

**A.**

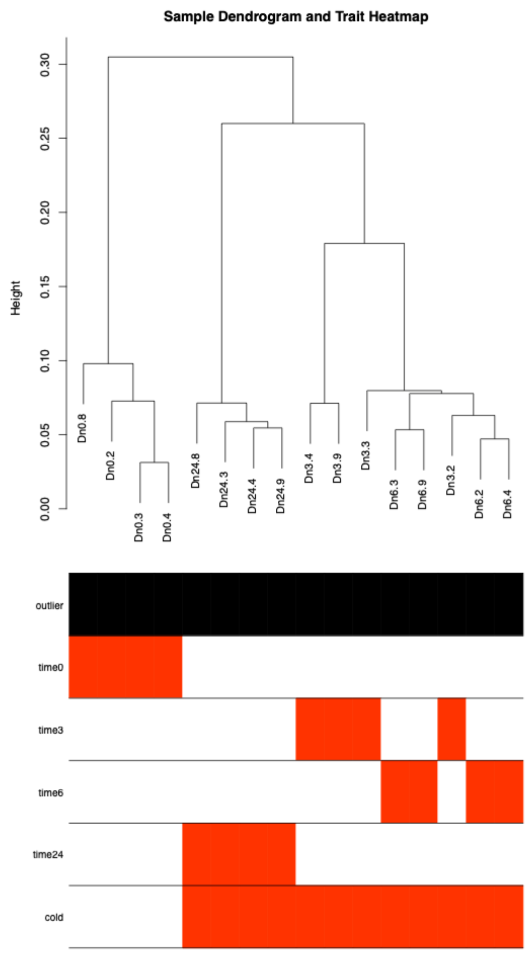


**B.**

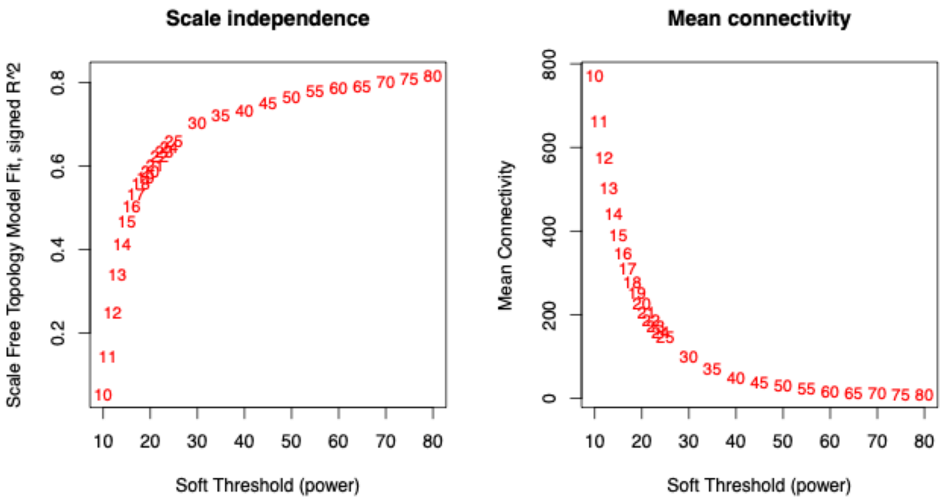


**Supplementary figure 3. Background information for *Draba nivalis* WGCNA.** A: Plot showing clustering of samples, B: Soft Thresholding Powers plotted against Scale Free Topology Fit (left) and Mean Connectivity (right).

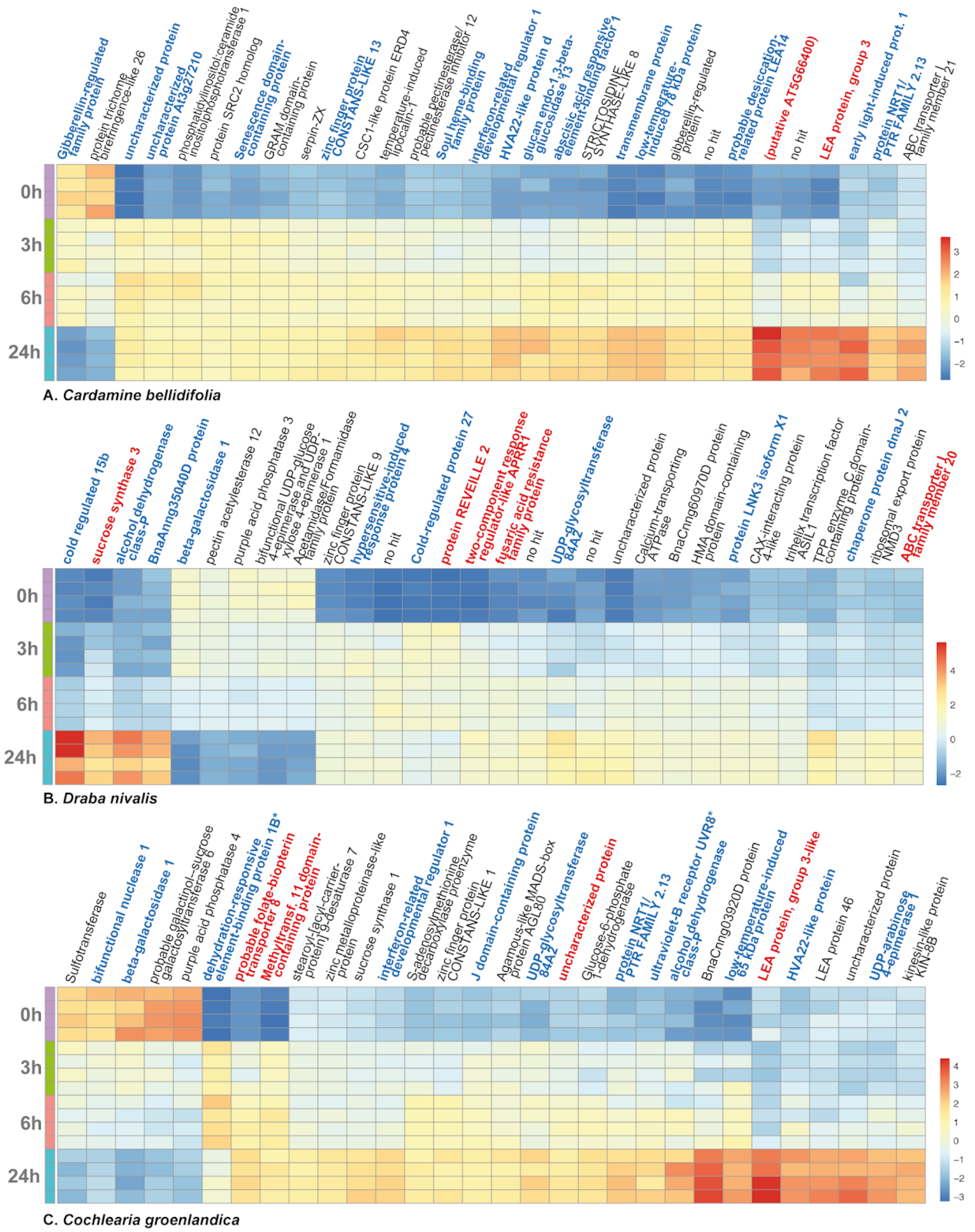


**Supplementary figure 4. Heatmaps of the most significantly differentially expressed genes (DEGs): A) *Cardamine bellidifolia*, B) *Draba nivalis*, and C) *Cochlearia groenlandica*.** Color scale = log2 transformed counts. Each row corresponds to a replicate, and there are four replicates at each time point (0h, 3h, 6h, 24h). Gene names in **bold and blue** = Found as 24h DEG in *Arabidopsis thaliana* and all Arctic species (based on *A. thaliana* homologs), gene names in **bold and red** = Found as 24h DEG only in Arctic species (based on *A. thaliana* homologs). *dehydration-responsive element-binding protein 1B = DREB1b/CBF1, ultraviolet-B receptor UVR8 = TCF1.

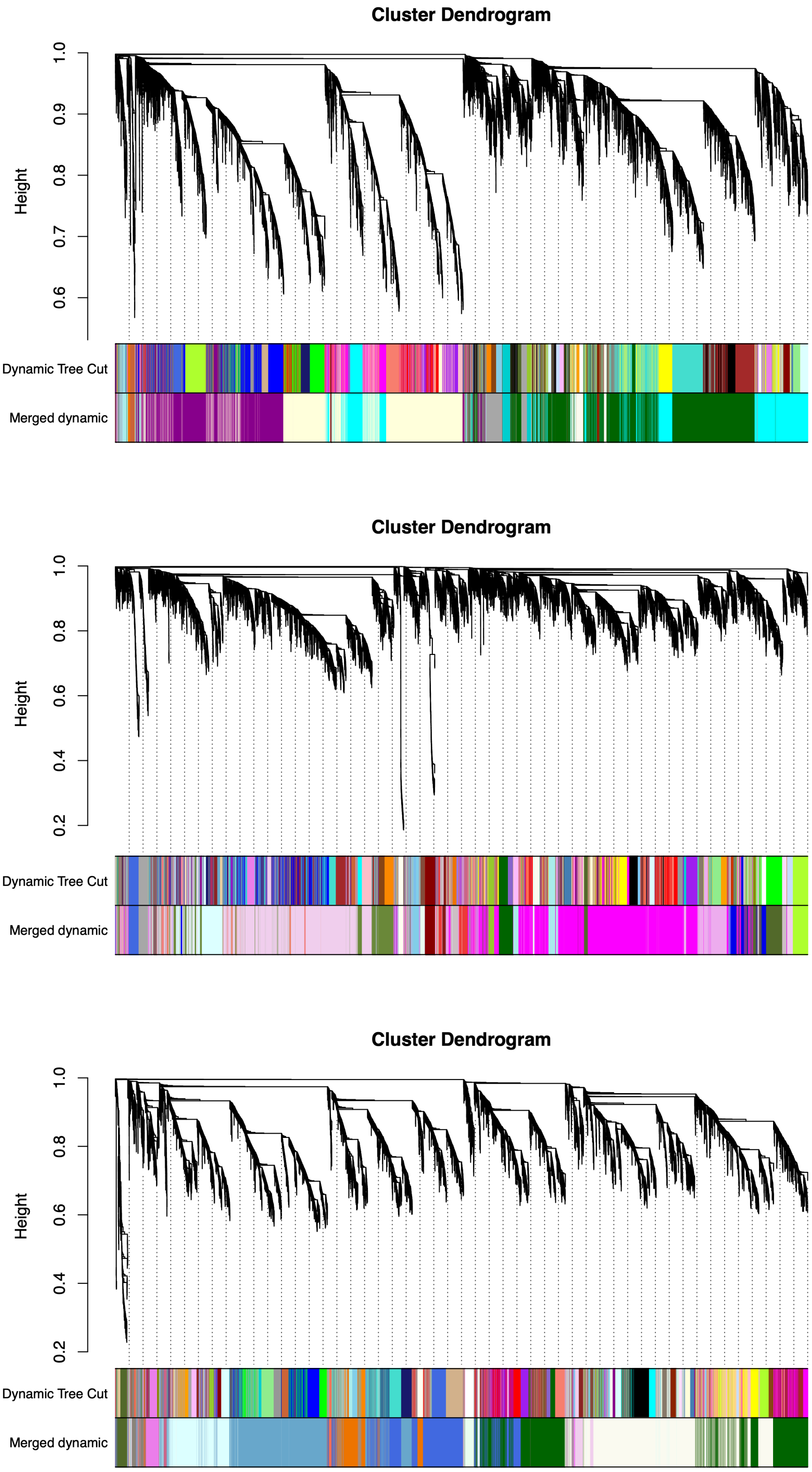


**Supplementary figure 5. Dendrograms with module colors (WGCNA).** Top: *Cardamine bellidifolia*, Middle: *Cochlearia groenlandica*, Bottom: *Draba nivalis*

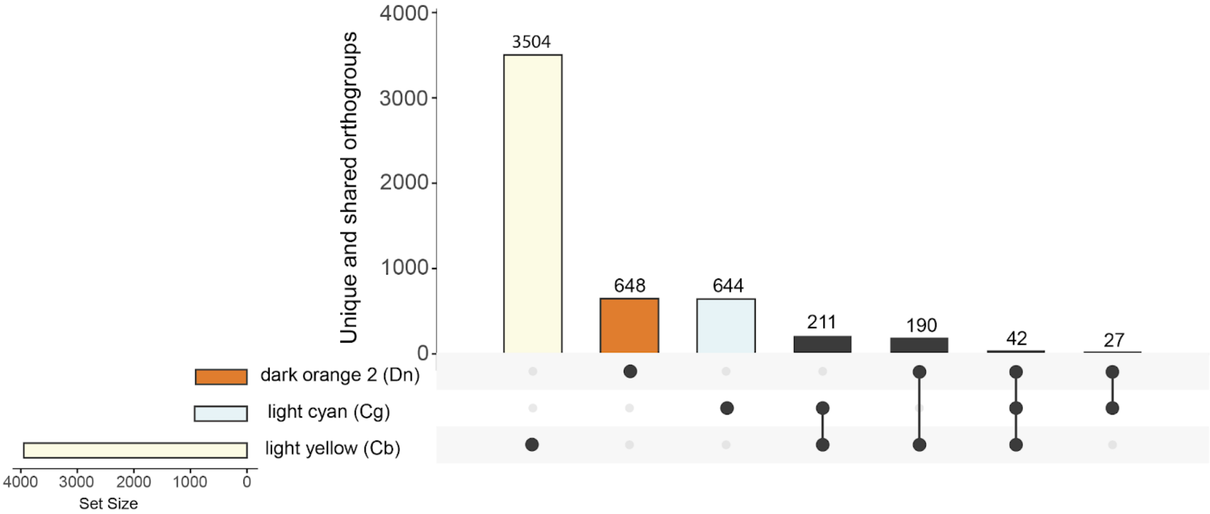


**Supplementary figure 6. UpSet plot of modules most positively correlated with the binary cold trait in each species.** The plot in the left corner shows total numbers of orthogroups, and the main plot shows the number of unique orthogroups, followed by the orthogroups that overlapped among modules (connected dots). Cb = *Cardamine bellidifolia*, Dn = *Draba nivalis*, Cg = *Cochlearia groenlandica.*

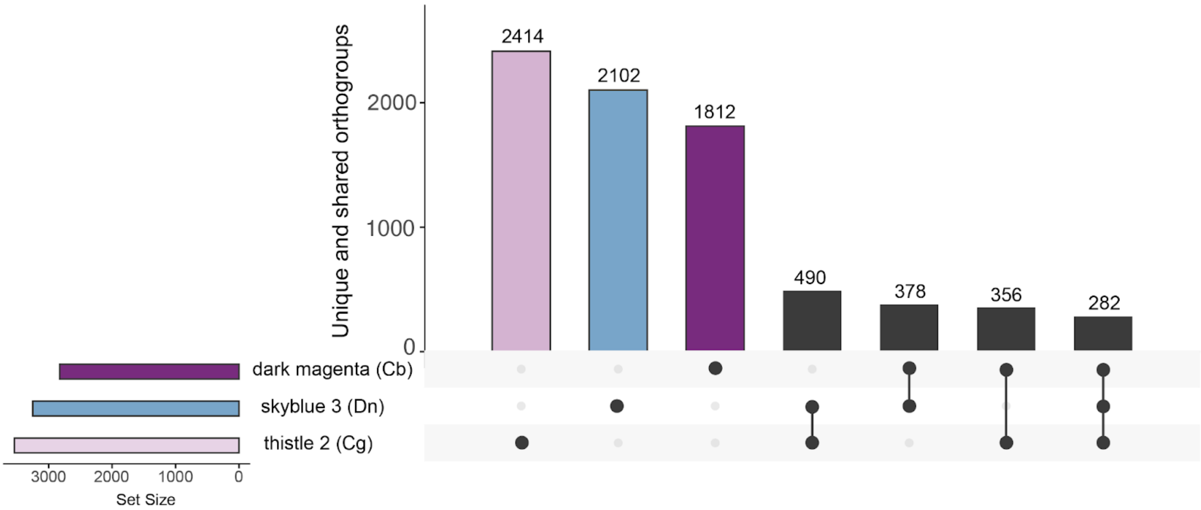


**Supplementary figure 7. UpSet plot of modules most positively correlated with 24h of cold in each species.** The plot in the left corner shows total numbers of orthogroups, and the main plot shows the number of unique orthogroups, followed by the orthogroups that overlapped among modules (connected dots). Cb = *Cardamine bellidifolia*, Dn = *Draba nivalis*, Cg = *Cochlearia groenlandica*

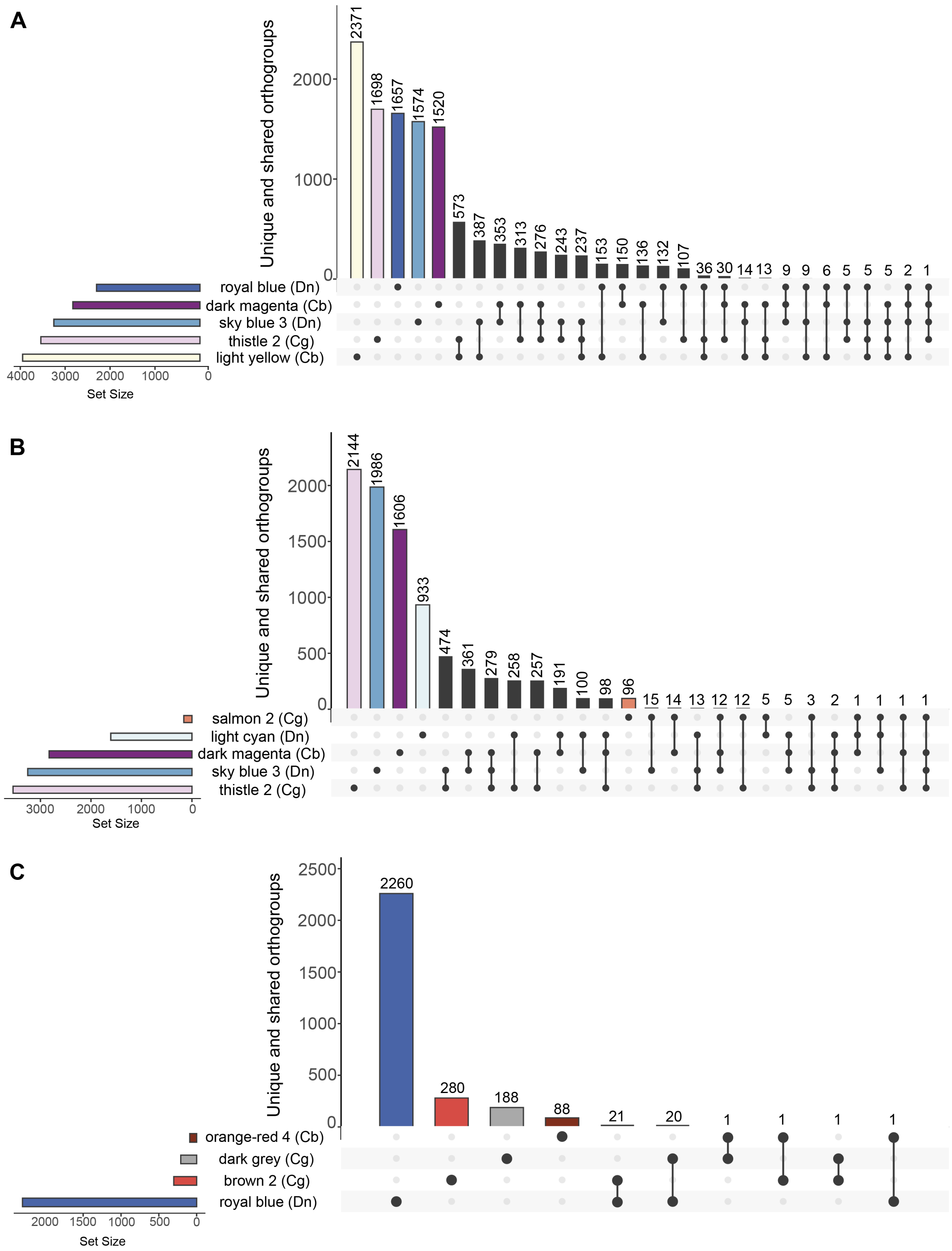


**Supplementary figure 8. UpSet plot of modules positively correlated with the binary cold trait, as well as 3h, 6h, 24h of cold (≥ 0.70).** The plot in the left corner shows total numbers of orthogroups, and the main plot shows the number of unique orthogroups, followed by the orthogroups that overlapped among modules (connected dots). **A:** All modules (showing only the largest modules/overlaps), **B:** All 24h modules, **C:** All 3h and 6h modules. Cb = *Cardamine bellidifolia*, Dn = *Draba nivalis*, Cg = *Cochlearia groenlandica.*

**Table S1. Sampling information for Arctic focal species**

| Species | Population ID | Region (Country) | Locality | Coordinates (DMS Latitude, DMS Longitude) |
|---|---|---|---|---|
| *Cardamine bellidifolia* | LG09-A-68-06-01 | Alaska (US) | Atigun Pass, Brooks Range | N68°8'10.44" W149°28'37.56" |
| *Cochlearia groenlandica* | LG09-A-48-09 | Alaska (US) | Nome, Seward Peninsula | N64° 29' 4.48" W165° 25' 49.9" |
| *Draba nivalis* | HHG-008-7-L-m | Alaska (US) | Waterfall Creek W | N63°2'42' W147°12'3.6" |

**Table S2. Overlapping DEGs among time points in *Cardamine bellidifolia***

A. Overlapping DEGs among time points.

| | Compared with 3h: | Compared with 6h | Compared with 24h |
|---|---|---|---|
| 3h shared [unique] | 1007 [0] | 760 [247] | 623 [384] |
| 6h shared [unique] | 760 [278] | 1038 [0] | 709 [329] |
| 24h shared [unique] | 623 [1887] | 709 [1801] | 2510 [0] |

B. Overlapping **upregulated** DEGs among time points.

| | Compared with 3h: | Compared with 6h | Compared with 24h |
|---|---|---|---|
| 3h shared [unique] | 855 [0] | 678 [177] | 518 [337] |
| 6h shared [unique] | 678 [194] | 872 [0] | 590 [282] |
| 24h shared [unique] | 518 [777] | 590 [705] | 1295 [0] |

C. Overlapping **downregulated** DEGs among time points.

| | Compared with 3h: | Compared with 6h | Compared with 24h |
|---|---|---|---|
| 3h shared [unique] | 152 [0] | 82 [70] | 97 [55] |
| 6h shared [unique] | 82 [84] | 166 [0] | 117 [49] |
| 24h shared [unique] | 97 [1118] | 117 [1098] | 1215 [0] |

**Table S3. Comparison of all DEGs among time points in *Cochlearia groenlandica***

A. Overlapping DEGs among time points.

| | Compared with 3h: | Compared with 6h | Compared with 24h |
|---|---|---|---|
| 3h shared [unique] | 729 [0] | 506 [233] | 516 [213] |
| 6h shared [unique] | 506 [497] | 1003 [0] | 685 [318] |
| 24h shared [unique] | 516 [2481] | 685 [2312] | 2997 [0] |

B. Overlapping **upregulated** DEGs among time points.

| | Compared with 3h: | Compared with 6h | Compared with 24h |
|---|---|---|---|
| 3h shared [unique] | 517 [0] | 369 [148] | 354 [163] |
| 6h shared [unique] | 369 [260] | 629 [0] | 448 [181] |
| 24h shared [unique] | 354 [1172] | 448 [1078] | 1526 [0] |

C. Overlapping **downregulated** DEGs among time points.

| | Compared with 3h: | Compared with 6h | Compared with 24h |
|---|---|---|---|
| 3h shared [unique] | 212 [0] | 137 [75] | 161 [51] |
| 6h shared [unique] | 137 [237] | 374 [0] | 237 [137] |
| 24h shared [unique] | 161 [1310] | 237 [1234] | 1471 [0] |

**Table S4. Comparison of all DEGs among time points in *Draba nivalis***

A. Overlapping DEGs among time points.

| | Compared with 3h: | Compared with 6h | Compared with 24h |
|---|---|---|---|
| 3h shared [unique] | 683 [0] | 606 [77] | 487 [196] |
| 6h shared [unique] | 606 [970] | 1576 [0] | 982 [594] |
| 24h shared [unique] | 487 [2341] | 982 [1846] | 2828 [0] |

B. Overlapping **upregulated** DEGs among time points.

| | Compared with 3h: | Compared with 6h | Compared with 24h |
|---|---|---|---|
| 3h shared [unique] | 501 [0] | 446 [55] | 349 [152] |
| 6h shared [unique] | 446 [548] | 994 [0] | 591 [403] |
| 24h shared [unique] | 349 [1130] | 591 [888] | 1479 [0] |

C. Overlapping **downregulated** DEGs among time points.

| | Compared with 3h: | Compared with 6h | Compared with 24h |
|---|---|---|---|
| 3h shared [unique] | 182 [0] | 160 [22] | 137 [45] |
| 6h shared [unique] | 160 [422] | 582 [0] | 389 [193] |
| 24h shared [unique] | 137 [1212] | 389 [960] | 1349 [0] |

**Table S8. Results from the supertest.** Significance of observed intersections between differentially expressed orthogroups in *Draba nivalis* (2167 differentially expressed orthogroups), *Cochlearia groenlandica* (2340 differentially expressed orthogroups), *Cardamine bellidifolia* (1986 differentially expressed orthogroups), and *Arabidopsis thaliana* (2428 differentially expressed orthogroups). Each orthogroup is counted only once per each species.

| Intersection | Degree | Background* | Observed overlap** | Expected overlap | Fold Enrichment | *p-value* |
|---|---|---|---|---|---|---|
| *C. groenlandica, D. nivalis* | 2 | 14,717 | 667 | 344.55 | 1.94 | 8.426537e-81 |
| *C. bellidifolia, D. nivalis* | 2 | 13,945 | 604 | 308.62 | 1.96 | 1.043355e-74 |
| *C. bellidifolia, C. groenlandica* | 2 | 14,052 | 635 | 330.72 | 1.92 | 1.435591e-160 |
| *C. bellidifolia A. thaliana* | 2 | 14,268 | 613 | 337.96 | 1.81 | 1.221776e-61 |
| *C. groenlandica A. thaliana* | 2 | 14,169 | 711 | 400.98 | 1.77 | 9.055851e-69 |
| *D. nivalis A. thaliana* | 2 | 14,182 | 768 | 371.00 | 2.07 | 2.261915e-114 |
| *C. groenlandica, D. nivalis, A. thaliana* | 3 | 12,684 | 367 | 76.53 | 4.80 | 5.176323e-150 |
| *C. bellidifolia, C. groenlandica, A. thaliana* | 3 | 12,702 | 334 | 69.94 | 4.78 | 8.524704e-135 |
| *C. bellidifolia, D. nivalis, A. thaliana* | 3 | 12,748 | 324 | 64.30 | 5.04 | 2.918931e-137 |
| ***C. bellidifolia, C. groenlandica, D. nivalis*** | **3** | **12,429** | **318** | **65.19** | **4.88** | **1.079363e-130** |
| *C. bellidifolia, C. groenlandica, D. nivalis, A. thaliana* | 4 | 11,715 | 212 | 15.21 | 13.94 | 4.218279e-173 |

*The background was set to the number of orthogroups shared between the species in each comparison. **Observed overlap includes orthogroups also shared with other species in the dataset (i.e. the observed overlap of 318 in *C. bellidifolia*, *C. groenlandica*, *D. nivalis*, includes the 212 orthogroups shared with *A. thaliana*).

**Table S18. Shared significantly enriched GO-terms** (fisher test in combination with the elim algorithm, $p < 0.05$) within the Biological Process (BP), Cellular Component (CC), and Molecular Function (MF) domains in Arctic species. The 24h DEG sets were also compared with those of *Arabidopsis thaliana* from Park et al. (2015), and **bold** indicates that the GO-term only was shared by Arctic species.

**A. Shared GO-terms in upregulated DEG sets**

| **Upregulated DEGs, 3h:** |
|---|
| GO:0080167 response to karrikin (BP) |
| GO:0009873 ethylene-activated signaling pathway (BP) |
| GO:0007623 circadian rhythm (BP) |
| GO:0006355 regulation of transcription, DNA-templated (BP) |
| GO:0009409 response to cold (BP) |
| GO:0009719 response to endogenous stimulus (BP) |
| GO:0010200 response to chitin (BP) |
| GO:0009631 cold acclimation (BP) |
| GO:0009414 response to water deprivation (BP) |
| GO:0009651 response to salt stress (BP) |
| GO:0031225 anchored component of membrane (CC) |
| GO:0005634 nucleus (CC) |
| GO:0080054 low-affinity nitrate transmembrane transporter activity (MF) |
| GO:0003677 DNA binding (MF) |
| GO:0003700 DNA-binding transcription factor activity (MF) |
| **Upregulated DEGs, 6h:** |
| GO:0008300 isoprenoid catabolic process (BP) |
| GO:0080167 response to karrikin (BP) |
| GO:0010501 RNA secondary structure unwinding (BP) |
| GO:0009873 ethylene-activated signaling pathway (BP) |
| GO:0007623 circadian rhythm (BP) |
| GO:0009737 response to abscisic acid (BP) |
| GO:0006355 regulation of transcription, DNA-templated (BP) |
| GO:0009409 response to cold (BP) |
| GO:0016115 terpenoid catabolic process (BP) |
| GO:0009719 response to endogenous stimulus (BP) |
| GO:0010378 temperature compensation of the circadian clock (BP) |
| GO:0009631 cold acclimation (BP) |
| GO:0009753 response to jasmonic acid (BP) |
| GO:0010017 red or far-red light signaling pathway (BP) |
| GO:0009414 response to water deprivation (BP) |
| GO:0048579 negative regulation of long-day photoperiodism, flowering (BP) |
| GO:0009651 response to salt stress (BP) |
| GO:0009269 response to desiccation (BP) |
| GO:0005634 nucleus (CC) |
| GO:0080054 low-affinity nitrate transmembrane transporter activity (MF) |
| GO:0003677 DNA binding (MF) |
| GO:0005509 calcium ion binding (MF) |
| GO:0003700 DNA-binding transcription factor activity (MF) |

**Upregulated DEGs, 24h:**

GO:0080167 response to karrikin (BP)
GO:0010501 RNA secondary structure unwinding (BP)
GO:0009873 ethylene-activated signaling pathway (BP)
GO:0007623 circadian rhythm (BP)
GO:0009737 response to abscisic acid (BP)
GO:0006355 regulation of transcription, DNA-templated (BP)
GO:0009409 response to cold (BP)
GO:0009408 response to heat (BP)
GO:0006121 mitochondrial electron transport, succinate to ubiquinone (BP)
GO:0009631 cold acclimation (BP)
GO:0009813 flavonoid biosynthetic process (BP)
GO:0006979 response to oxidative stress (BP)
GO:0009414 response to water deprivation (BP)
GO:0009651 response to salt stress (BP)
GO:0051555 flavonol biosynthetic process (BP)
GO:0010286 heat acclimation (BP)
GO:0010224 response to UV-B (BP)
**GO:0009636 response to toxic substance (BP)**
**GO:0008295 spermidine biosynthetic process (BP)**
**GO:0019319 hexose biosynthetic process (BP)**
GO:0008378 galactosyltransferase activity (MF)
GO:0003700 DNA-binding transcription factor activity (MF)
GO:0035251 UDP-glucosyltransferase activity (MF)
**GO:0003677 DNA binding (MF)**
**GO:0008177 succinate dehydrogenase (ubiquinone) activity (MF)**
**GO:0008792 arginine decarboxylase activity (MF)**

**B. Shared GO-terms in downregulated DEG sets**

**Downregulated DEGs, 3h:**

GO:0048046 apoplast (CC)

**Downregulated DEGs, 6h:**

GO:0009416 response to light stimulus (BP)
GO:0055088 lipid homeostasis (BP)

**Downregulated DEGs, 24h:**

GO:0006949 syncytium formation (BP)
GO:0009638 phototropism (BP)
GO:0009828 plant-type cell wall loosening (BP)
**GO:0009639 response to red or far red light (BP)**
**GO:0010112 regulation of systemic acquired resistance (BP)**
**GO:0009734 auxin-activated signaling pathway (BP)**
**GO:0009624 response to nematode (BP)**
**GO:0045490 pectin catabolic process (BP)**
GO:0005576 extracellular region (CC)
GO:0009505 plant-type cell wall (CC)
**GO:0031225 anchored component of membrane (CC)**
**GO:0016021 integral component of membrane (CC)**
GO:0030570 pectate lyase activity (MF)

**Table S24. 24h DEGs under positive selection based on Birkeland *et al.,* (2020).**
PSG = Positively selected gene

| | Total number of PSGs Birkeland et al. (2020) | PSGs among all DEGs (3h, 6h, 24h) | PSGs among upregulated DEGs (3h, 6h, 24h) | PSGs among downregulated DEGs (3h, 6h, 24h) |
|---|---|---|---|---|
| ***C. bellidifolia*** | 201 | 40 | 20 | 20 |
| ***C. groenlandica*** | 159 | 25 | 12 | 13 |
| ***D. nivalis*** | 360 | 65 | 36 | 29 |

**Table S25. 24h DEGs with convergent substitutions based on Birkeland *et al.,* (2020).**
Cb = *Cardamine bellidifolia* 24h DEG set, Cg = *Cochlearia groenlandica* 24h DEG set, Dn = *Draba nivalis* 24h DEG set

| | Total number of convergent genes Birkeland et al. (2020) | Convergent genes among all DEGs (3h, 6h, 24h) | Convergent genes among upregulated DEGs (3h, 6h, 24h) | Convergent genes among downregulated DEGs (3h, 6h, 24h) |
|---|---|---|---|---|
| ***C. bellidifolia*, *C. groenlandica*** | 58 | 14 Cb<br>12 Cg | 9 Cb<br>10 Cg | 5 Cb<br>2 Cg |
| ***C. groenlandica, D. nivalis*** | 33 | 8 Cg<br>9 Dn | 6 Cg<br>3 Dn | 2 Cg<br>6 Dn |
| ***D. nivalis*, *C. bellidifolia*** | 126 | 23 Dn<br>23 Cb | 16 Dn<br>18 Cb | 7 Dn<br>5 Cb |

**Table S29. Gene Ontology terms that were overrepresented in the light yellow (*C. bellidifolia*) light cyan (*C. groenlandica*) and dark orange 2 (*D. nivalis*) co-expression modules. These modules were the most positively correlated with the binary measure of cold in each species.**

| GO Identifier | GO Term Name | Domain |
|---|---|---|
| GO:0080167 | response to karrikin | BP |
| GO:0007623 | circadian rhythm | BP |
| GO:0009737 | response to abscisic acid | BP |
| GO:0006355 | regulation of transcription, DNA-templated | BP |
| GO:0009409 | response to cold | BP |
| GO:0009719 | response to endogenous stimulus | BP |
| GO:0009753 | response to jasmonic acid | BP |
| GO:0097305 | response to alcohol | BP |
| GO:0010017 | red or far-red light signaling pathway | BP |
| GO:0009414 | response to water deprivation | BP |
| GO:0009651 | response to salt stress | BP |
| GO:0005634 | nucleus | CC |
| GO:0003677 | DNA binding | MF |
| GO:0005509 | calcium ion binding | MF |
| GO:0043565 | sequence-specific DNA binding | MF |
| GO:0003700 | DNA-binding transcription factor activity | MF |

**Table S42. Gene Ontology terms that were overrepresented in the dark magenta (*C. bellidifolia*) thistle 2 (*C. groenlandica*) and sky blue 3 (*D. nivalis*) co-expression modules. These modules were the most positively correlated with 24h with cold in each species.**

| GO Identifier | GO Term Name | Domain |
|---|---|---|
| GO:0009790 | embryo development | BP |
| GO:0042254 | ribosome biogenesis | BP |
| GO:0046686 | response to cadmium ion | BP |
| GO:0009553 | embryo sac development | BP |
| GO:0006413 | translational initiation | BP |
| GO:0046034 | ATP metabolic process | BP |
| GO:0006364 | rRNA processing | BP |
| GO:0005654 | nucleoplasm | CC |
| GO:0005743 | mitochondrial inner membrane | CC |
| GO:0044451 | obsolete nucleoplasm part | CC |
| GO:0000502 | proteasome complex | CC |
| GO:0031461 | cullin-RING ubiquitin ligase complex | CC |
| GO:0005730 | nucleolus | CC |
| GO:0009506 | plasmodesma | CC |
| GO:0032040 | small-subunit processome | CC |
| GO:0005829 | cytosol | CC |
| GO:0005634 | nucleus | CC |
| GO:0005774 | vacuolar membrane | CC |
| GO:0008026 | helicase activity | MF |
| GO:0003743 | translation initiation factor activity | MF |

**Supplementary Text 1. Note on positively selected/convergent cold-responsive genes**
We found several cold responsive genes that have previously been identifyed to be under positive selection or contain convergent substitutions in the three Arctic species (identified in Birkeland et al. 2020; Mol. Biol. Evol.). This included *COR15B* and *CSDP1* in *D. nivalis*, *LEA4-5* in *C. groenlandica*, and a highly upregulated transmembrane protein (putative homolog of *A. thaliana* AT1G16850) in *C. bellidifolia*. We found that a gene with convergent substitutions in all Arctic species, *EMB2742*, was upregulated in all species (and in *A. thaliana*). Some convergent genes showed different expression patterns depending on the species. For instance, *CAT2* was downregulated in *D. nivalis* and upregulated in *C. groenlandica*. This gene has previously been found to contain convergent substitutions in *D. nivalis* and *C. groenlandica*, and to be under positive selection in *D. nivalis*. The low temperature responsive transcription factor *RAV1* has previously been found to be under positive selection in *C. groenlandica*, but was not differentially expressed in this species. We also note that *MAPKKK14*, (previously found to contain convergent substitutions in the Arctic species) was upregulated in all species.
